# Distribution of Gene Tree Topologies with Duplication, Loss, and Coalescence

**DOI:** 10.64898/2026.01.19.700405

**Authors:** Sarthak R. Mishra, Matthew W. Hahn

## Abstract

Both duplication and loss can generate gene trees that differ from the species tree, and consequently many methods can be used to infer the number and timing of gene duplication and loss events from comparisons of gene trees and species trees. However, most such reconciliation methods use a model of gene duplication that does not include the coalescent process, or that restricts it in important ways. As a result, changes to tree topologies due to coalescence will lead to extra duplications and losses using these methods, events that did not actually occur. Here, we present results from the multispecies coalescent with duplication and loss (MSC-DL) model, which allows for the unrestricted interaction between duplication, loss, and coalescence. Theoretical results show that even histories with only a single duplication event can lead to many more trees than are considered without accounting for coalescence: for a species tree with 2 tips, 9 trees are possible, while with 6 tips, more than 19 million trees are possible; adding even a single loss almost doubles the number of possible topologies. We also calculate the probabilities of different gene tree topologies and their branch lengths under the MSC-DL for cases with two species and one duplication. These results have important implications for the accuracy of reconciliation methods, ortholog identification methods, and our understanding of evolutionary histories of duplication and loss.

## Introduction

Evolutionary biology relies heavily on phylogenetic trees to infer the history of organisms. There are two types of phylogenetic trees that are widely used: gene trees, which represent the history of individual loci, and species trees, which represent the broader evolutionary relationships among species (Maddison [1997]). Understanding the evolution of gene trees within species trees is essential for understanding the evolutionary history of genes and organisms.

However, it is well known that the topology of a gene tree may not match the corresponding species tree topology—a phenomenon called gene tree discordance (Maddison [1997], Degnan and Rosenberg [2009]). There are several biological factors that can cause gene tree discordance, including incomplete lineage sorting, duplication, loss, introgression, and horizontal gene transfer; we largely consider the first three here. Incomplete lineage sorting (ILS) occurs when two or more lineages do not coalesce in a shared population, but instead coalesce in an ancestral population that contains additional lineages (Maddison [1997]). In the context of the multispecies coalescent (MSC) model (Hudson [1983], Rannala and Yang 2003]), ILS occurs when the time between two speciation events is short, and can lead to the generation of discordant gene trees. Gene duplication increases the number of lineages in a gene tree, generating gene tree branches that do not exist in the species tree, and trees that are therefore discordant. Loss decreases the number of lineages in a gene tree, leading to further discrepancies between gene trees and species trees (Albalat and Cañestro [2016]).

Reconciliation methods represent a wide and varied set of algorithms that attempt to infer the evolutionary events necessary to explain any discordance between gene trees and species trees (reviewed in Boussau and Scornavacca [2020], Williams et al. [2024]). Some algorithms assume that discordance is only due to duplication and loss (so-called “DL” models; e.g. Goodman et al. [1979], Page and Charleston [1997], Guigo et al. [1996], Chen et al. [2000], Zmasek and Eddy [2001], Durand et al. [2006]), while some additionally consider the contribution of horizontal gene transfer (“DTL” models; e.g. Hallett and Lagergren [2001], Bansal et al. [2012], Stolzer et al. [2012], Szöllősi et al. [2013], Morel et al. [2020]). Importantly, ILS can cause discordance on its own, and ILS (and coalescence more generally), duplication, and loss can interact with each other, making the problem of gene tree reconciliation even more complicated. Thus, several reconciliation methods have been introduced that allow for the interaction of duplication, loss, and coalescence (Rasmussen and Kellis [2012], Wu et al. [2014], Paszek et al. [2021], Mishra et al. [2024]).

Every reconciliation algorithm has an underlying generative model, even if it is only implicit. These models describe the ways in which evolutionary events such as duplication, loss, or coalescence can change a gene tree, and therefore which and how many events the algorithm will use to reconcile the gene tree with the species tree. DL and DTL models do not include coalescence, and therefore all coalescence events leading to discordance will be interpreted by these reconciliation methods as extra duplications and losses (Hahn [2007]). While the earliest models of duplication, loss, and coalescence only allowed certain types of coalescence (Rasmussen and Kellis [2012]), several recent models have been proposed that allow for unrestricted coalescence (Li et al. [2021], Mishra et al. [2024], Li et al. [2024]).

Our goal in this paper is to examine the properties of one of these generative models, the multispecies coalescent model with duplication and loss (MSC-DL; Mishra et al. [2024]). This model allows duplication, loss, and coalescence to interact without restriction, and can therefore provide the clearest picture to date of the frequency and topology of gene trees produced by the interaction of these events. Such results will be of great help in designing, interpreting, and understanding the limitations of reconciliation algorithms (and related tasks) that are dependent on identifying duplication and loss events.

In this work we:

1. Formalize the space of gene tree topologies generated under the MSC-DL model.
2. Derive a closed-form formula for the number of distinct gene tree topologies generated from a single duplication (coupled with coalescence) depending on the number of species in a tree, as well as the number of topologies from one duplication, one loss, and coalescence.
3. Compute exact probability distributions for all gene tree topologies in the two-species case with one duplication, as well as branch lengths in this case.

## The MSC-DL Framework

### A brief description of the MSC-DL model

The multispecies coalescent with duplication and loss (MSC-DL) model is a unified model of gene family evolution that allows gene duplication, gene loss, and coalescence. The MSC-DL is an extension of the standard multispecies coalescent model (MSC) that further allows duplication, loss, and coalescence between gene copies. A full description of the MSC-DL can be found in Mishra et al. [2024], but here we describe details that are most salient to our analysis.

The MSC-DL treats each individual locus as evolving via the MSC, where discordance between individual gene trees and the species tree can only arise via ILS. Each duplication event inserts a DNA sequence from one locus—termed the parent locus—into a pre-existing locus—called a daughter locus. We say that the duplication *event* itself happens at the parent locus (red star in Figure 1a), while the duplicated sequence *inserts* at the daughter locus (arrow in Figure 1b). Because the daughter locus existed in the genome prior to the duplication event (though it was not the “daughter” locus at that point), this locus has its own gene tree drawn from the MSC. The gene tree at the daughter locus exists regardless of the duplication event or timing, and it is not changed by the insertion of the duplicated DNA; the duplicated sequence simply follows the daughter tree from the point of insertion. If multiple gene tree lineages exist at the daughter locus at the time of duplication, one is chosen randomly for the insertion. Depending on the recombination distance between the parent and daughter loci, their topologies and branch lengths may not be exactly the same. In what follows we will assume that the loci are completely recombinationally unlinked, but this assumption is easily relaxed. Loss occurs at a single time on a single branch, and can occur either at a parent locus (whether or not there has been a duplication) or a daughter locus (when there has been a duplication). All lineages descending from the loss event are removed from the corresponding gene tree.

**Fig. 1.**
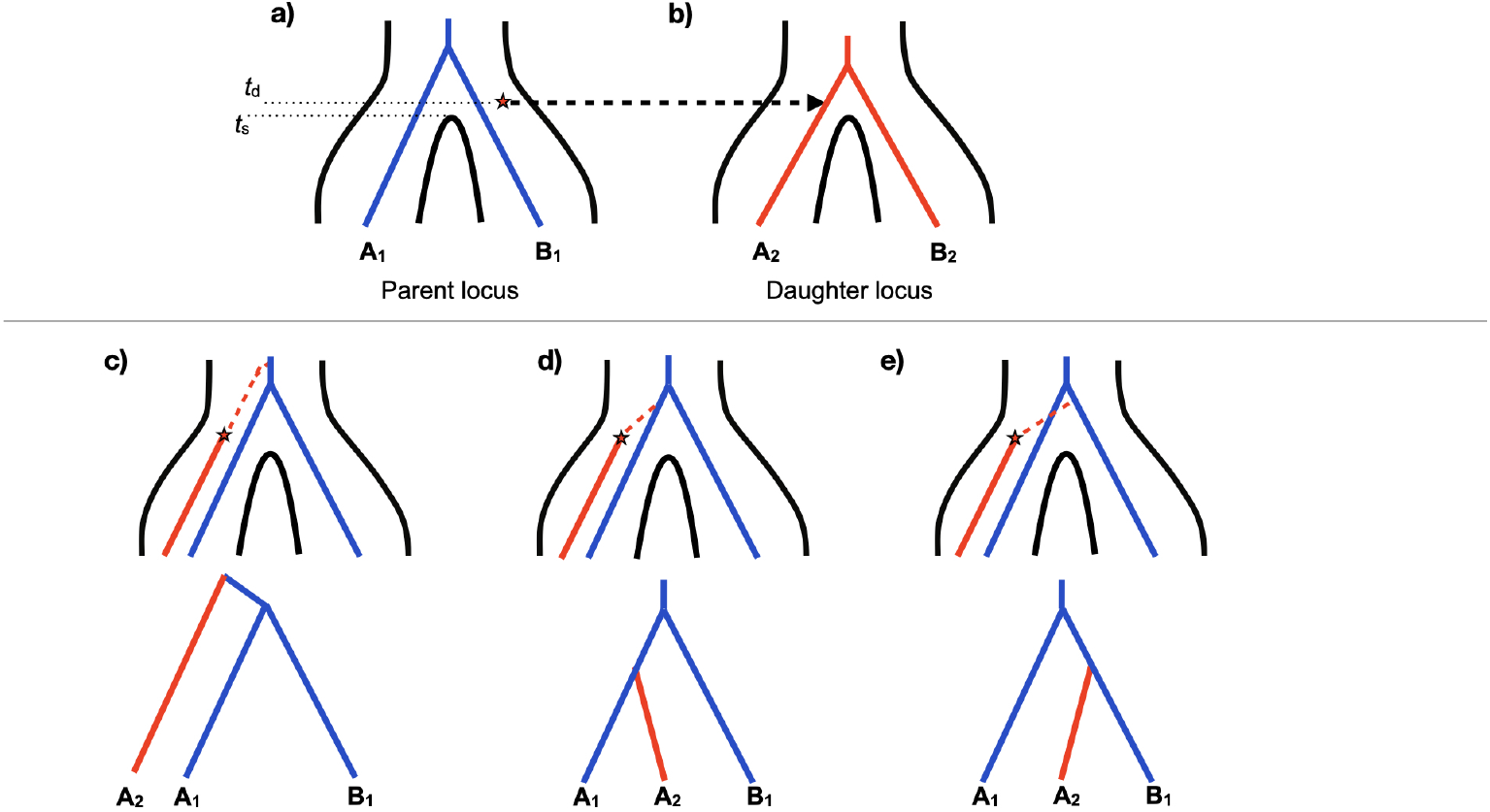
An example from the MSC-DL. In all panels, a two-species tree with speciation time, *t_s_*, is shown. **a)** A gene tree at the parent locus (blue) is derived from the MSC. A duplication event (red star) occurs at this locus at time *t_d_*. **b)** A gene tree at the daughter locus (red), which is also drawn from the MSC. The duplication is inserted at the daughter locus at the point where the arrow meets the gene tree, which must also occur at time *t_d_*. **c-e)** The MSC-DL describes the possible ways that the parent and daughter loci coalesce to form the full gene tree. In all panels the star and the red branch come from the daughter gene tree, while the blue branches come from the parent gene tree. The coalescent process that attaches the the two trees is shown as a red dashed branch above the star. The trees in the bottom row are simply unfolded views of the resulting gene trees.

In addition to a coalescent process acting independently at the parent and daughter loci, the daughter locus also must coalesce with the parent locus after a duplication event (dashed branch in Figure 1c-e). This process occurs because the duplicated sequence was itself a lineage that existed at the parent locus before the duplication event, with a genealogical history at that locus that it carries with it to the daughter locus (see, e.g., Figure S2 in Mishra et al. [2024]). At the time of duplication, *t_d_*, the MSC-DL allows the daughter copy to freely coalesce with any branch of the parent locus that comes after the time of duplication (with time moving backward from the present) according to a standard MSC process. We refer to the lineage at the parent locus that it coalesces with as the *attachment* branch. Importantly, this attachment does not have to occur immediately, and coalescence between the loci can lead to additional discordance.

Thus, the MSC-DL model involves three coalescent processes: 1) coalescence at the parent locus, 2) coalescence at the daughter locus, and 3) coalescence between the parent and daughter loci. Given these processes, the MSC-DL model defines a probability distribution over gene tree topologies given a fixed species tree.

### Example calculation in the MSC-DL

Figure 1b shows the interaction of duplication with coalescence at the daughter locus in determining the insertion branch, while panels c, d, and e show the interaction of the parent gene tree and the coalescent process in determining the attachment branch. In Figure 1a,b, the duplication occurs and inserts at *t_d_*, after the time of speciation between species *A* and *B, t_s_*. We usually assume that the mutational process that creates duplicates is Poisson-distributed along the parent gene tree branches, but for now we condition on a single duplication event and will therefore ignore the stochasticity introduced by the mutation process.

Even though the duplication has occurred in the ancestral population of species *A* and *B*, it does not necessarily insert into a branch of the gene tree at the daughter locus that includes both species. In the example shown, *t_d_* occurs before coalescence in the daughter locus gene tree (again, looking backward in time), the probability of which corresponds to the probability of the daughter lineage not coalescing before the time of duplication, which is given by 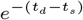. In this case, the duplication still has two lineages into which it can insert, *A* or *B*, with equal probability. Hence, for a duplication that occurs after speciation, the probability that it lands on either tip branch of the daughter gene tree is given by 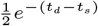 for each branch.

Once the duplication inserts into a daughter gene tree lineage (in Figure 1 it is lineage *A*), it still has to coalesce with the gene tree at the parent locus. Figures 1c,d,e show the three different trees that can be produced by this single duplication, though these three trees are not equal in probability. To see why this is, let us denote samples from the parent locus with the subscript “1” and from the daughter locus with the subscript “2.” The most common topology caused by this duplication will be (*A*_2_, (*A*_1_, *B*_1_)), with the lineages from the parent tree coalescing together first. This is most common because these lineages have probability of coalescing of 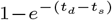 after the speciation event but before the duplication event. When this occurs, the resulting gene tree topology must be (*A*_2_, (*A*_1_, *B*_1_)). If the lineages from the parent locus have not coalesced by *t_d_* (which happens with probability 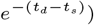, then all three topologies are equiprobable, each with frequency 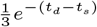. Together, the probability of the tree (*A*_2_, (*A*_1_, *B*_1_)), assuming the insertion shown, is 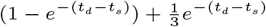 (Figure 1).

There are a few things to note about this example. First, conditioning on a duplication event that happens at a different time (e.g. before speciation) will give different probabilities for both the insertion branch and the attachment branch, and we use the example here simply to show the types of calculations that must occur. Second, regardless of when duplication occurs, insertion and attachment are unrestricted under the MSC-DL (*contra* earlier models; Rasmussen and Kellis [2012]). This means that every gene tree branch that can arise under the MSC has a non-zero probability of serving as either an insertion branch or an attachment branch, though attachment branches must exist after the time of duplication. Third, in the specific example shown in Figure 1, ILS *per se* does not occur, since all events are happening in the same ancestral population; nonetheless, coalescence is generating the different possible gene trees. The results in the following sections follow directly from the properties of the MSC-DL laid out here.

## Results

### Explosion of gene tree space with one duplication

The MSC-DL requires coalescence of duplicate lineages inserted at the daughter locus with lineages in the parent locus. This process increases the number of possible gene trees (“gene tree space”) significantly. To show the size of the gene tree space, we start by explaining the process with just one duplication.

#### Number of duplication branches

When a duplication occurs, it must insert into a specific branch of the gene tree at the daughter locus (Figures 1 and 2); we call this a “duplication branch.” Without conditioning on any particular time of duplication, the number of lineages where a duplication can insert is the total number of unique lineages (branches) at the daughter locus that are possible given the species tree and the coalescent process. The number of duplication branches can also be expressed as the summation of all possible trees for all possible combinations of lineages:

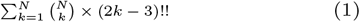

where *k* ∈ *{*1, …, *N}*, (2*k −* 3)!! denotes the number of rooted binary trees for *k* taxa in a sub-tree and *N* denotes the total number of species. This equation accounts for all the *k* lineages that can possibly make a clade from a tree with *N* species and multiplies it with all possible binary trees with *k* lineages. Here is an example for *N* = 3:

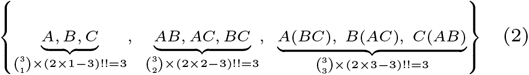

**Fig. 2.**
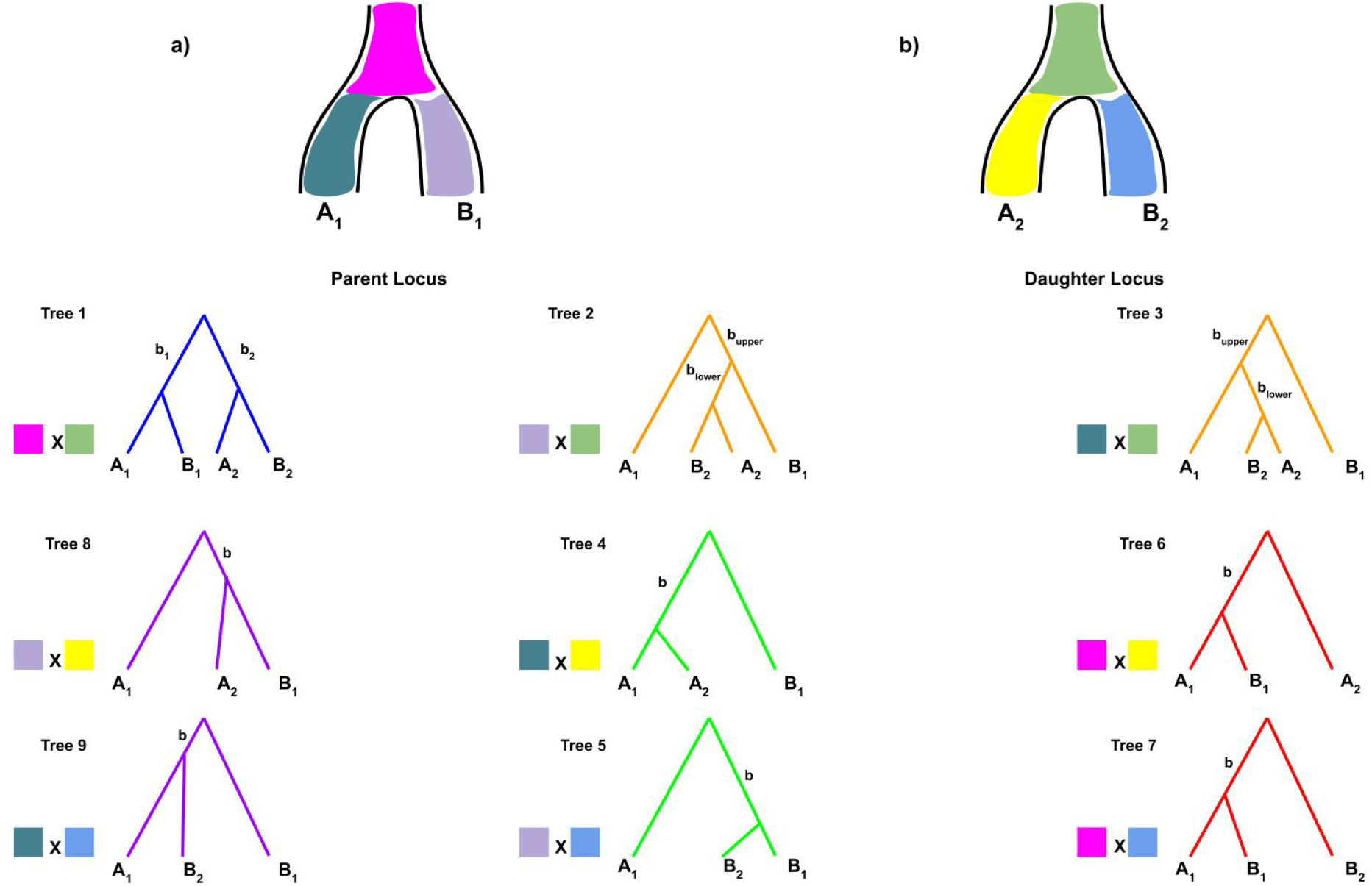
Enumerating all possible gene trees in the two-species case. a) Possible lineages where the daughter gene tree can attach to the parent gene tree ((A,B), A, or B). Although the branches of the species tree are colored, these correspond to branches of gene trees with the same identities. **b)** Possible branches that a duplicate can be inserted into on the daughter gene tree ((A,B),A, or B). **First row** (trees 1,2,3): Three different gene trees with insertion on lineage (A,B) and attachment with ((A,B), A, or B). **Second row** (trees 8,4,6): Three different gene trees with insertion in lineage A and attachment with ((A,B), A, or B). **Third row** (trees 9,5,7): Three different gene trees with insertion in lineage B and attachment with ((A,B), A, or B). **All rows**: Gene trees with the same color branches have the same probability expressions and are represented as the same color in other figures and tables.

In this example, the first term describes the case in which a duplicate can land on a branch that extends to one of the three tip lineages (*A, B, C*). The second part of the formula accounts for all possible tree topologies in which the duplicate can land on an internal branch of the tree, with the three possible clades subtended by this branch shown (*AB, AC, BC*). Finally, a duplicate can land on the branch subtending all three taxa, and there are three possible topologies for such a tree. The formula is therefore calculating all branches that subtend from 1 to *N* lineages, along with the possible tree topologies associated with each number of lineages. For *N* =3, all of this shows that there are 9 possible gene tree branches on which a duplicate can insert at the daughter locus. (For 2 species, there are 3 possible gene tree branches on which a duplicate can insert see Figure 2].)

#### Number of attachment branches

In addition to characterizing all the lineages where duplication can insert into, we must also enumerate all the places where the duplicated lineage can coalesce (i.e. attach) back to on the parent gene tree (Figures 1 and 2). The duplicated lineage can attach to the parent gene tree on all branches from all parent topologies drawn from the MSC that are older than (above) the time of duplication. However, it should also be noted that each unique attachment branch is associated with an entire tree topology. For instance, attachment on the *A*_1_ branch of the parent tree must be associated with all topologies containing *A*_1_, of which there are three in the three-species case. Hence, the possible attachment branches include all the branches for all the gene trees possible with MSC, which is always equal to or greater than the number of branches that a duplicate can insert on.

We can express the total number of branches that the daughter gene tree can attach to in the parent gene tree as the product of:

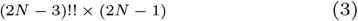

where *N* denotes the number of species, (2*N −* 3)!! denotes the number of rooted binary trees for *N* taxa, and (2*N −*1) denotes the number of branches in a rooted binary tree for *N* taxa. For a three-species tree, this implies that there are 15 unique attachment branches. (For 2 species, there are only 3 unique attachment branches see Figure 2].)

#### Total number of gene tree topologies with one duplication

The total number of gene tree topologies possible with just one duplication in an *N* -species tree is the product of the number branches on which a duplication can insert into on the daughter gene tree and the number of branches on which it can attach to on the parent tree. This formula is:

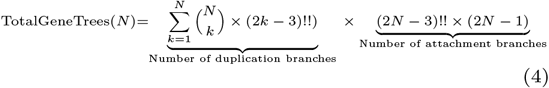

More details can be found in Supplementary section 1. By comparison, the number of gene trees possible without duplication or loss (i.e. due to the MSC alone) for a species tree with *N* species is given by the number of attachment branches for a tree with *N −* 1 species (see Table 1).

**Table 1.** Number of gene trees possible with just 1 duplication and coalescence, as a function of the number of species, *N*.

| $N$ | Possible Duplication Branches | Possible Attachment Branches | Total Trees |
| --- | --- | --- | --- |
| 2 | 3 | 3 | 9 |
| 3 | 9 | 15 | 135 |
| 4 | 37 | 105 | 3885 |
| 5 | 225 | 945 | 212625 |
| 6 | 1881 | 10395 | 19552995 |

Figure 2 shows the result with one duplication graphically for a species tree with *N* = 2. There are three branches of possible gene trees at the daughter locus on which a duplicate can insert. Likewise, there are three branches of possible gene trees at the parent locus that a duplicated lineage can attach to. The product of all these possibilities results in the nine gene trees shown. For reference, in the standard DL model, it is assumed that one duplicate in a two-species tree can only produce three rooted trees: ((*A*_1_, *A*_2_), *B*_1_), ((*B*_1_, *B*_2_), *A*_1_), and ((*A*_1_, *A*_2_), (*B*_1_, *B*_2_)). Any tree that looks different from these must invoke more than a single duplication.

In Table 1, we show that the number of trees grows factorially as the number of species increases. For *N* = 2 species there are 9 possible gene trees possible with 1 duplication, for *N* = 3 there are 135 trees possible, and for *N* = 6 there are more than 19 million possible trees. Obviously, adding more duplication events will further complicate the situation, both by increasing the number lineages where duplication can occur and where a duplicate can attach. While we do not consider these cases here, in the next section we do consider the case of one duplication and one loss.

### Extension to one duplication and one loss

Adding one loss with one duplication increases the complexity of the problem. Losses increase the number of gene trees by creating “extra” lineages and gene trees—those that are not possible without loss. For example, in a two-species tree, the gene tree (*A*_1_, *A*_2_) cannot be generated with one duplication and no losses. In this section, we present a mathematical expression for the number of additional trees with one duplication (1D), one loss (1L), and coalescence.

#### Extra attachment branches

When we model the interaction of duplication and loss, we first have to determine the order of events (e.g. did the duplication happen before loss or vice versa). If the loss happened further in the past than the duplication, then the lost lineage must be at the parent locus—otherwise, there is no duplicated sequence at the daughter locus to lose. If a loss is more recent than the duplication, the loss can occur either on a parent gene tree or daughter gene tree.

Interestingly, the trees produced when a single loss lands on the daughter gene tree are all already possible in the 1D case—i.e. any loss on a daughter gene tree can be produced with 1 duplication and coalescence (Supplementary section 2.1). There is one exception: when the loss lands on the root of the daughter lineage, the result is complete loss of the daughter copy. In this case the resulting gene tree is just the parent gene tree by itself, which cannot be produced with 1D+coalescence.

A more interesting case occurs when the loss lands on the parent gene tree. These resulting trees are not possible with just one duplication. Therefore, to understand the added complexity from one duplication and one loss, we must examine the consequence of losses that occur at the parental locus; specifically, we want to count the additional attachment branches created by such a loss. With a single loss, the parental gene tree can lose from 1 to *N −*1 number of leaves (we ignore the case in which the entire parent locus is lost). A loss on a branch subtending *M* leaves will result in a parental tree with *k* = *N − M* tips remaining. This results in new attachment branches (in new trees), the number of which are also given by equation 3, substituting *k* for *N*.

#### Total number of gene tree topologies with one duplication and one loss

Because losses at the daughter locus can also be produced by 1D+coalescence, all of these trees are also produced by 1D+1L. Therefore, the total number of gene tree topologies produced with one duplication and one loss is equal to all of the trees produced in the 1D case (equation 4) plus all of the extra gene trees produced by losses at the parent locus.

To calculate the additional gene trees that can be produced by 1D+1L, we must multiply all the extra attachment branches (previous sub-section) by the possible duplication branches (which does not change relative to the 1D case). The total number of additional trees is therefore given by the sum total of all these effects:

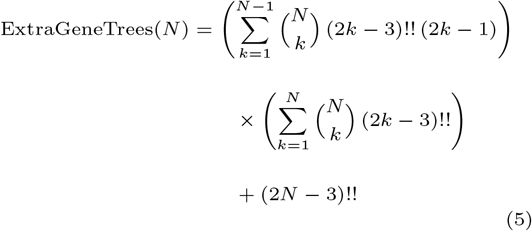

The added term at the end represents the case when the entire daughter copy is lost, which results in a gene tree including only tips at the parent locus (producing an additional (2*N −* 3)!! trees). Note, however, that these calculations do not count duplication on the root of the parent tree followed by loss of the parent tree on the same branch, as this case would leave only the daughter tree; including these topologies would add additional trees. Further details can be found in Supplementary section 2.

Finally, the total number of gene tree topologies in the 1D+1L case is the sum of equations 4 and 5 for any particular *N*. In Supplementary Table 1 we show the number of additional trees that are produced by one duplication, one loss, and coalescence. Summing these numbers and the ones in Table 1 shows that there are 16 topologies in the *N* = 2 case and more than 33 million topologies for *N* = 6. Together, our results demonstrate that just one duplication and one loss can almost double the possible gene trees compared to the one-duplication case.

### Gene tree probabilities in the two-species case

Although many gene trees are possible under the MSC-DL, not all topologies will occur with equal probability. In fact, many of the trees included in Table 1 may not be found in nature, as their probability is very low. This can be seen by considering the fact that some marginal trees (i.e. those at either the parent or daughter loci) are quite rare under the MSC even without the added complexity of duplication and loss. In this section we explore the probability distribution of gene trees under the MSC-DL. Enumerating all possible gene tree probabilities is difficult, especially as the number of species (and consequently the number of gene trees) grows. Here, we explicitly show these calculations for one particular gene tree topology in a two-species tree; all gene tree probabilities for the two-species case with one duplication are shown in Table 2.

**Table 2.**
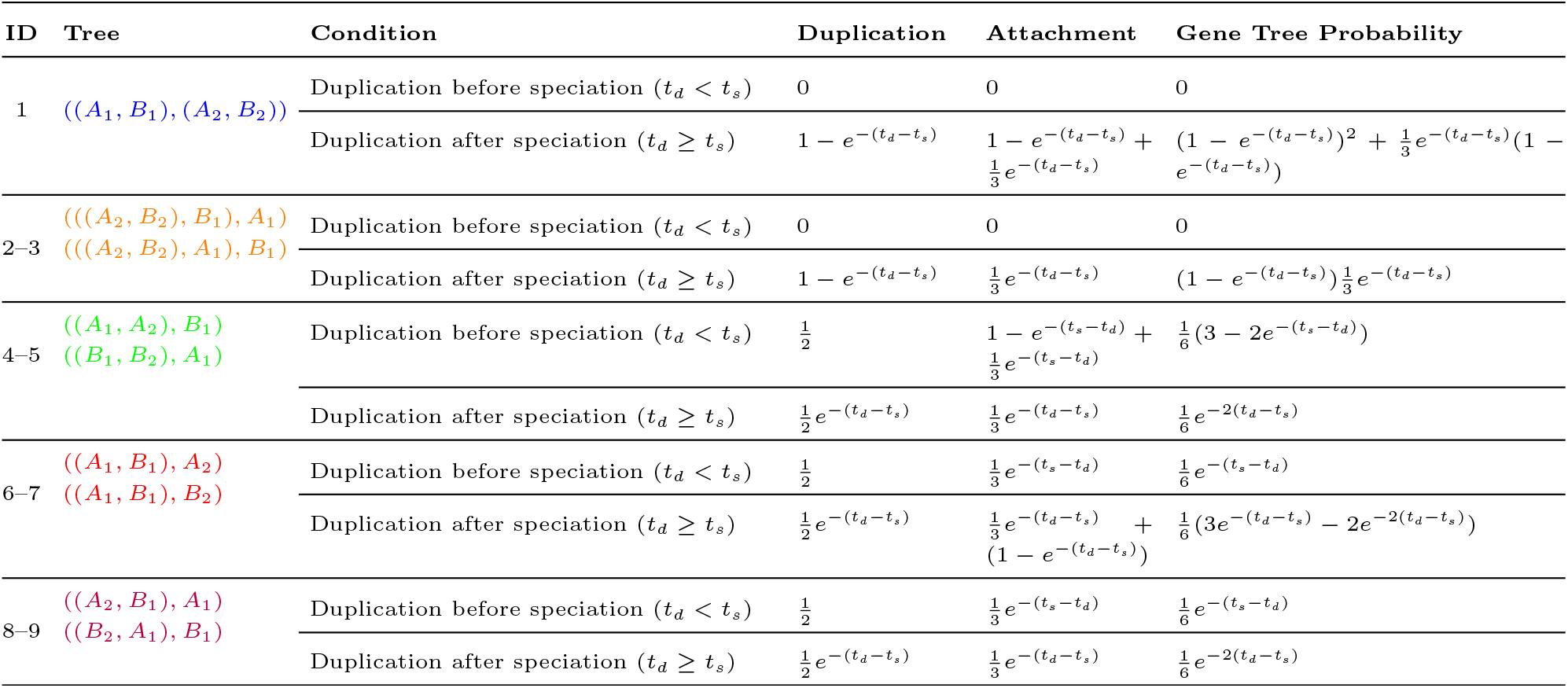
The probability of different gene tree topologies, including duplication insertion and attachment probabilities, in a two-species tree with one duplication.

For a given time of duplication, the probability of a gene tree is calculated by determining which branch the duplicate is inserted into at the daughter locus and which branch the duplicated lineage attaches to at the parent locus. For instance, to obtain a gene tree of the form (*A*_2_, (*A*_1_, *B*_1_)) (tree 6) with only a single duplication, the duplication must insert into lineage *A* at the daughter locus, followed by an attachment to the branch subtending (*A*_1_, *B*_1_) at the parent locus. Moreover, the duplication in this example can occur either before or after the time of speciation (looking backwards in time), depending on coalescence at the daughter locus (Supplementary Figure 2). Thus, to compute the probability of a gene tree, conditional on a time of duplication, we must express all the possible ways a duplicate can insert onto a specific branch at the daughter locus and all the possible ways the duplicated lineage can attach to a specific branch at the parent locus.

For duplication before speciation (*t_d_ < t_s_*), the probability that the duplicate inserts onto the *A* branch (which is necessary to produce tip *A*_2_) is 1*/*2 since there are only two branches possible. Given this insertion, in order to form the tree (*A*_2_, (*A*_1_, *B*_1_)) the daughter branch must not coalesce with the parent branch before speciation. This occurs with probability 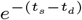. If it has not coalesced, there are three different possible topologies that can form when all three descendant lineages make it back to the ancestral population, each of which has probability 1*/*3. Therefore, the probability of this tree topology given *t_d_ < t_s_* is:

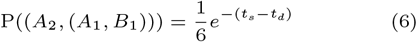

For the same tree with duplication after (or at the same time as) speciation (*t_d_ ≥ t_s_*), things are slightly more complex. In this case, there must be no coalescence at the daughter locus between the *A* and *B* lineages (which occurs with probability 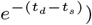, and the duplicate must insert on the *A* branch (with probability 1*/*2). At the parent locus, we can form this topology one-third of the time if the parent gene tree has not coalesced (with probability 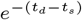) and every time it has (with probability 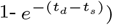. The total probability of this tree topology given *t_d_ ≥ t_s_* is then:

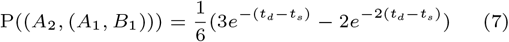

In Table 2 we give the probabilities for all 9 possible gene trees when there are 2 species and 1 duplication. For each tree we list the probabilities of duplication and attachment separately, as well as the total gene tree probability (which is just the product of the duplication and attachment terms), and separate probabilities for the distinct regimes of duplication before and after speciation. While every tree is possible when the duplication occurs after speciation, trees with (*A*_2_, *B*_2_) pendant branches cannot have duplication occurring before speciation.

These probabilities are also shown graphically in Figure 3 across different times of duplication; note that the different gene tree probabilities sum to 1 at any given time of duplication. The colors for the expression values in the figure and table match the color scheme in Figure 2: if two trees have the same probability expression, then they are colored the same. Despite having 9 distinct topologies, we only see 5 distinct probability expressions because of symmetries between the *A* and *B* lineages.

**Fig. 3.**
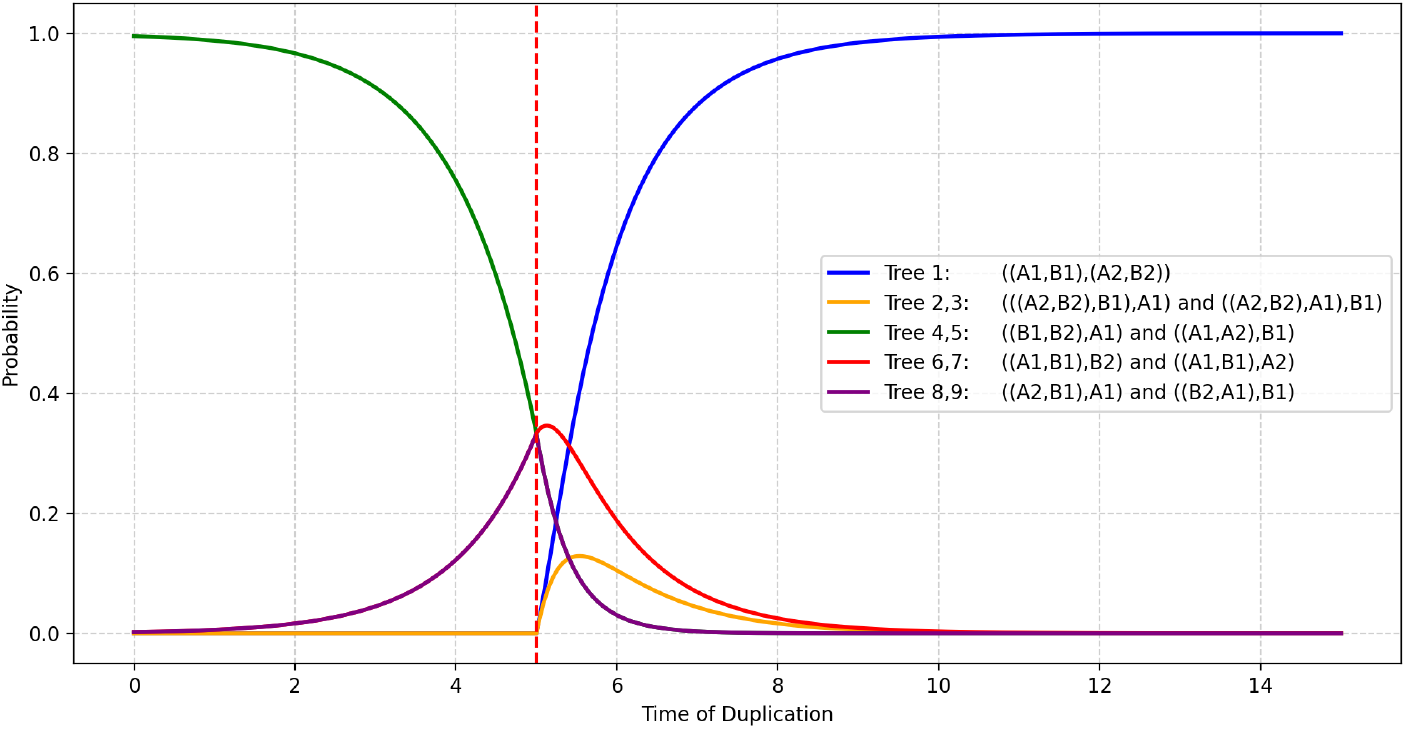
The relative probability of gene trees varies with the time of duplication, *t_d_* (x-axis), relative to the time of speciation, *t_s_* (vertical dashed line). At the time of speciation (*t_s_*=5) trees 6,7, 4,5, and 8,9 are all equiprobable at 0.33. All results come from Table 2. Time in the figure is measured in coalescent units.

One interesting observation about the probabilities in Figure 3 is that they converge to the standard DL results when duplication occurs long before or long after speciation. That is, with *t_d_ << t_s_* (i.e. a duplication on a tip branch), we are highly likely to see either ((*A*_1_, *A*_2_),*B*_1_) or ((*B*_1_, *B*_2_),*A*_1_), because coalescence has occured between parent and daughter long before speciation. Likewise, when *t_d_ >> t_s_* (i.e. a duplication on the shared ancestral branch), we are highly likely to see ((*A*_1_, *B*_1_),(*A*_2_, *B*_2_)) because coalescence at both the parent and daughter locus has occurred long before duplication. It is only close to the speciation time—both before and after—that coalescence generates many more topologies (in this two-species case in which there can be no discordance at either marginal locus).

### Expected branch lengths in the two-species case

In addition to the probabilities of individual gene tree topologies, it would be useful to have the expected branch lengths for each tree. Therefore, we found the expected branch lengths for all gene trees with 2 species and 1 duplication as a function of *t_d_* and *t_s_*.

A complication arises when finding expected branch lengths while conditioning on a specific number of duplication events (and not a specific time of duplication). When there is a non-zero duplication rate (*λ*) per unit time, the height of the marginal gene tree at the parent locus directly affects the number of duplications observed. For instance, taller trees will lead to more duplications. As a result, the branch length distributions for trees with one duplication may not match familiar theoretical expectations at the limit *λ* → 0.

Below, we derive the distribution of heights of the gene trees as a function of *λ*. To clarify the effect of conditioning on a number of duplications, in the main text we only show the calculations for the tree with zero duplications, (*A*_1_, *B*_1_) (“tree 0”). Full derivations for all other trees are given in Supplementary section 3, but follow directly from the approach laid out here. Table 3 shows the internal branch lengths for all nine trees with one duplication.

**Table 3.** Expected internal branch lengths of trees in Figure 2. *λ* represents the duplication rate.

| Tree | Branch | Expectation | Equation (Supp) |
| --- | --- | --- | --- |
| Tree 1 | $b_1$ | 1 | 24 |
| Tree 1 | $b_2$ | $1 + \frac{1}{1+3\lambda} + \frac{1}{3+2\lambda}$ | 27 |
| Trees 2–3 | $b_{\text{upper}}$ | $\frac{1}{1+2\lambda}$ | 42 |
| Trees 2–3 | $b_{\text{lower}}$ | $\frac{1}{1+3\lambda} + \frac{1}{3+2\lambda}$ | 44 |
| Trees 4–5 | $b$ | $\mathbb{E}[b \mid \mathcal{E}_4] = \tilde{P}_1 \cdot \mathbb{E}[b \mid \mathcal{R}_1] + \tilde{P}_2 \cdot \mathbb{E}[b \mid \mathcal{R}_2] + \tilde{P}_3 \cdot \mathbb{E}[b \mid \mathcal{R}_3]$ | 70 |
| Trees 6–7 | $b$ | 1 | 84 |
| Trees 8–9 | $b$ | $\frac{1}{1+2\lambda}$ | 100 |

#### *No-duplication Tree* (*A*_1_, *B*_1_)

Given species tree (*A, B*) with speciation time *t_s_* and duplication rate *λ*, let *t_j_* denote the height of the parent gene tree (*A*_1_, *B*_1_). Our goal is to calculate the expected height, *E*[*t_j_* | (*A*_1_, *B*_1_)], for a given *λ* (here and below we assume that there is no loss). For this tree to be generated, zero duplications can occur on the parent tree (*n* = 0). We define this event as:

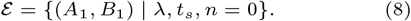

##### Prior distribution of the gene tree height

For two lineages, the waiting time until coalescence is exponentially distributed with rate 1. Because *A*_1_ and *B*_1_ must coalesce above speciation,

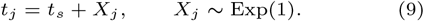

Hence,

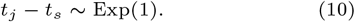

Thus, the density of *t_j_* is

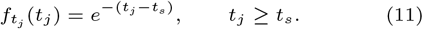

This is the standard result for the height of a gene tree under the coalescent model, assuming that there is no probability of duplication.

##### Conditional distribution of the duplication count

However, when there is duplication, to get this tree we must condition on zero duplications having occurred. The parent gene tree has two branches, each of length *t_j_*. Therefore the total branch length on which duplications may occur is 2*t_j_*

We assume that duplications occur according to a Poisson process with rate *λ*. Hence, the number of duplications, *n*, that fall on this tree is given by:

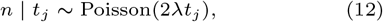

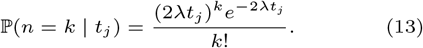

We want the probability for cases in which no duplications occur on the parent tree, i.e. *k* = 0:

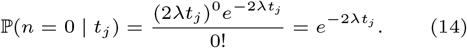

Since *E* can only occur when there is no duplication, this is also the probability of this event:

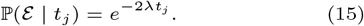

##### Posterior distribution

Putting together the prior distribution and the conditional distribution we can get the posterior distribution of tree height, *t_j_*. By Bayes’ rule,

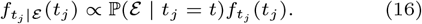

Substituting, we get:

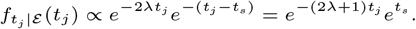

Since *t_s_* is constant, this simplifies to:

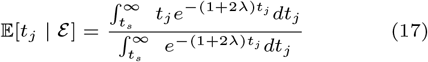

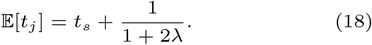

This equation provides the expected height for the parent tree on which no duplications have occurred, given a time of speciation and a duplication rate.

When duplication is impossible (*λ →* 0), the classic result is that the expected height of this gene tree is *t_s_* plus one coalescent unit (compare equations 9 and 18). For instance, with *t_s_* = 2, *t_j_* is expected to be 3. How does the result differ when *λ >* 0? For the same scenario with *λ* = 0.1, the expected height of this gene tree evaluates to:

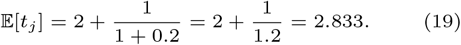

We can see that trees with 0 duplications are expected to be slightly shorter, as longer trees will accumulate more duplications.

Table 3 shows the expected internal branch lengths for all nine gene trees with one duplication (see Supplementary Section 3 for full derivations), with branches labeled as in Figure 2. Figure 4 shows these branch lengths as a function of *λ*, comparing the expected values to simulations carried out using the software *dupcoal* (Mishra et al., [2024). As can be seen, the match between theory and simulation is very good.

**Fig. 4.**
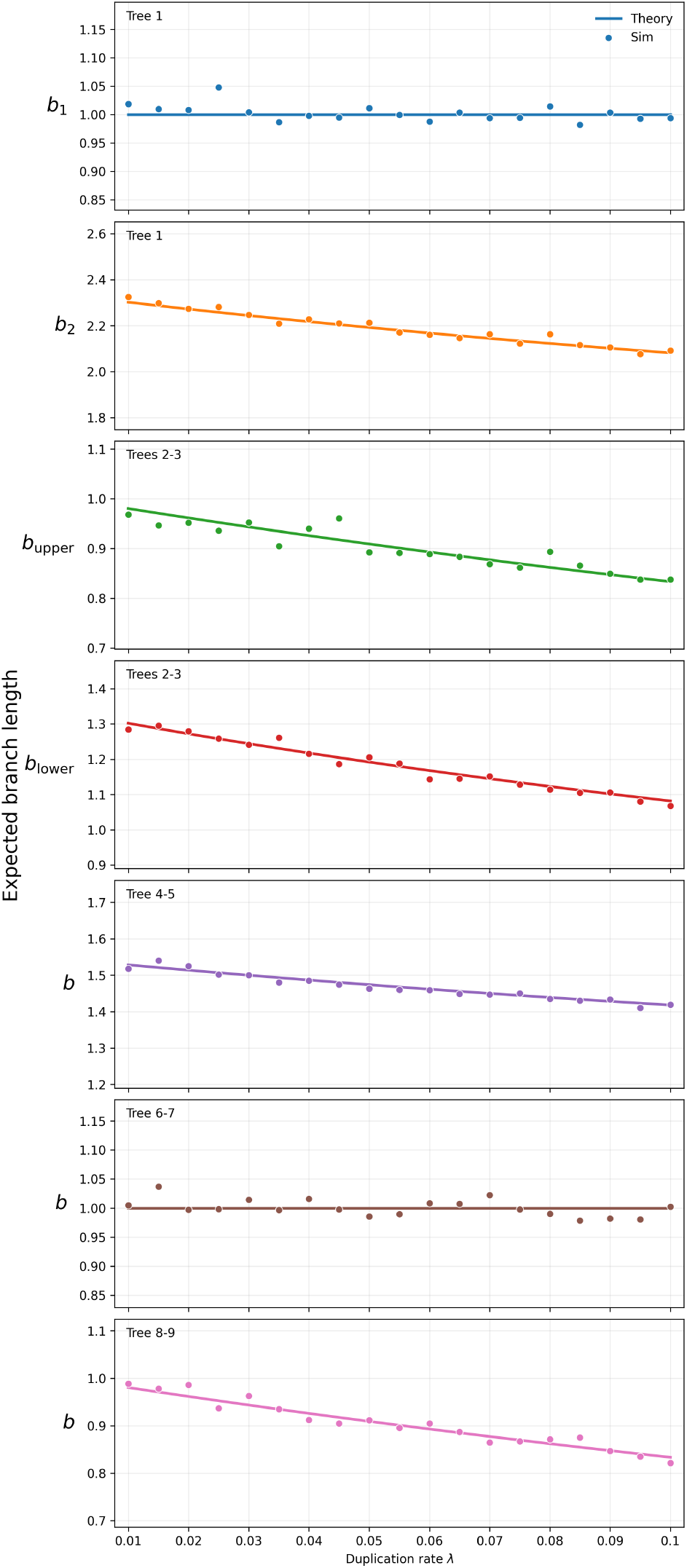
Expected branch lengths of internal branches on gene trees with one duplication. Trees and internal branches are labeled as in Figure 2, with lengths (y-axis) given in coalescent units. Theoretical expectations come from Table 3. Simulated data come from 500,000 gene trees generated with *dupcoal* (Mishra et al., [2024]) for each value of *λ*, with the average branch length for each topology among these trees shown.

## Discussion

In this work, we have explored the frequency and topology of gene trees under a unified model of duplication, loss, and coalescence. We show that gene tree space grows factorially with the number of species in a tree, with even a single duplication event leading to a huge number of possible gene trees. The biological processes we model are not unusual, nor do we consider extreme portions of parameter space. We have carried out many of our example calculations with trees containing only two species, both because these calculations are easier and to make the important point that these effects will be seen in even the simplest scenarios. With two species there can be no discordance of the marginal gene trees, and indeed ILS is not needed to produce discordance from the interactions of duplication and coalescence.

One of the main goals of our work is to demonstrate how ignoring the coalescent process can lead to many incorrect inferences about the numbers of duplications and losses occurring in evolutionary history, as well as their timing. Standard DL and DTL reconciliation methods do not consider coalescence, forcing coalescent events to be interpreted as additional duplications and losses. As mentioned, the model underlying DL reconciliation (either implicitly or explicitly) assumes that a single duplication in a two-species tree can only produce three distinct topologies, whereas we have shown that nine are possible. As an example of this mismatch, for trees 2,3 in

Figure 2, standard DL reconciliation will infer a history with two duplications and two losses. As species trees with more taxa are considered, more extreme incorrect inferences become likely; Supplementary Figure 3 shows a simple three-species tree with a single duplication and coalescence that will be inferred to have four duplications and eight losses by DL reconciliation. Based on the results presented here, we strongly recommend that researchers use reconciliation methods that allow for the interaction of duplication, loss, and coalescence (Rasmussen and Kellis [2012], Wu et al. [2014], Paszek et al. [2021], Mishra et al. [2024]).

In addition to inferring the number of duplications and losses, gene trees are often reconciled in order to identify orthologs (e.g. Emms and Kelly [2019], Zhang et al. [2020]). The general idea is that reconciliation can label nodes in a gene tree as either “duplication” or “speciation,” and by using trees containing only speciation nodes we can ensure that we are using orthologous genes for downstream tasks. However, the results presented here imply that these methods will identify the wrong nodes as speciation nodes (for example in trees 8,9), or simply miss speciation nodes. The underlying reason for this problem is that duplication events are always associated with coalescence events, and it is these latter events that form the nodes of gene trees, not the duplications themselves (star in Figure 1). While we know of no perfect solution to this problem (but see (Mishra et al. [2024] for one approach), it is fortunate that identifying speciation nodes may not be necessary for accurate species tree inference (Smith and Hahn [2021], Yan et al. [2022]).

Finally, we recognize that the results presented here are limited to very simple biological scenarios (i.e. one duplication), whereas real histories can be much more complex. Unfortunately, even in the simple scenarios considered, many types of inferences appear to be difficult or impossible. For example, even straightforward questions such as “Did this duplication occur before or after speciation” or “Does the history of this gene tree include a loss” do not seem answerable in all cases, at least when only topologies are used. We hope that future probabilistic approaches taking advantage of expected branch lengths may make it feasible to distinguish among a much larger set of histories. Alternatively, the huge size of gene tree space could make model-based probabilistic inference intractable, but that machine learning approaches would be able to use the large tree space and disparate types of data (topologies, branch lengths) to make accurate inferences.

## Author contributions statement

S.R.M. and M.W.H. conceived the paper, S.R.M. conducted the analyses, S.R.M. and M.W.H. interpreted the results. S.R.M. and M.W.H. wrote the manuscript.

## Acknowledgments

This work is supported in part by funds from the National Science Foundation (DBI-2146866).

