## Supplement pdf for "Distribution of Gene Tree Topologies with Duplication, Loss, and Coalescence"

### Supplementary Materials for "Distribution of Gene Tree Topologies with Duplication, Loss, and Coalescence"

#### 1 Insertion and Attachment in the 1 Duplication Case

##### 1.1 Number of possible branches duplicates can insert into (Daughter locus)

Let $N$ be the number of species. Under the MSC-DL model a duplication insertion can occur on any gene-tree branch that subtends a non-empty subset of taxa. We here show that the total number of such branches is:

$$\text{Possible Duplication Branches}(N) = \sum_{k=1}^{N} \binom{N}{k} (2k-3)!! \tag{1}$$

###### 1.1.1 Branches being counted

A duplication insertion can occur on all branch with non zero probability under the MSC-DL. Hence, we are counting all possible branches in all the gene tree topologies across all epochs of time in the species tree.

###### 1.1.2 All size subsets can appear in a clade

Under the multispecies coalescent (MSC) model, any subset of taxa can appear at any given time before the most recent common ancestor (MRCA) of a gene tree with non-zero probability (Degnan and Salter 2005). That means a branch where the duplication on a species tree landed can represent any subset of taxa below it. The number of such subsets is:

$$\sum_{k=1}^{N} \binom{N}{k} \tag{2}$$

###### 1.1.3 Shape of clades for each subset

For any fixed subset of size $k$ under the MSC, we can generate $(2k-3)!!$ number of rooted binary gene tree topologies. In all of these trees there is exactly one branch subtending these clades whose descendant leaves correspond to the chosen subset. The number of such trees is:

$$(2k-3)!! \tag{3}$$

###### 1.1.4 No double counting

Two different subsets of leaves (in section 1.1.2) correspond to different descendant sets of lineage. Hence, they have different and distinct associated branches. Within each subset of lineages each binary tree topology counts exactly one branch of the set in that shape, making equation 1 unique.

###### 1.1.5 Final expression

Summing over all subsets of lineage of all sizes and all the corresponding trees (clade shapes) that can be formed from them:

$$\text{Possible Duplication Branches}(N) = \sum_{k=1}^{N} \underbrace{\binom{N}{k}}_{\text{subsets of lineages}} \cdot \underbrace{(2k-3)!!}_{\text{shapes contributing one branch}} \tag{4}$$

#### 1.2 Number of possible branches duplicated lineages can attach to (Parent locus)

##### 1.2.1 Number of possible attachment branches

We now show that the number of possible branches where the duplicated lineage can attach in the parent locus is:

$$\text{Possible Attachment Branches}(N) = (2N-3)!!\,(2N-1) \tag{5}$$

##### 1.2.2 MSC generates all gene tree topologies

Under the MSC all rooted binary gene tree topology on sets of taxa occurs with non-zero probability. The number of such gene trees for $N$ species is:

$$(2N-3)!! \tag{6}$$

##### 1.2.3 Each topology contains branches

A rooted binary tree with $N$ leaves has exactly $(2N-1)$ branches.

##### 1.2.4 All branches can serve as an attachment point

Given a duplication event at time $t_d$, the duplicated lineage can coalesce with all branch of a parent locus with non-zero probability. This is because:

1. Under the MSC all topologies occurs with non-zero probability.
2. All branches in that topology exists for some time interval ($> t_d$) with non-zero probability.
3. The MSC-DL does not imposes any restrictions on coalescence between loci.
4. Hence, all branches in all MSC-possible topologies are a valid attachment location.

##### 1.2.5 No double counting

Two different topologies represent two unique set of branches. Given a topology, all branches are also unique. This makes lineages being counted by equation 5 unique.

##### 1.2.6 Final expression

The total number of branches in the daughter locus where a duplicate can attach is all possible branches in all possible gene trees .

$$\text{Possible Attachment Branches}(N) = \underbrace{(2N-3)!!}_{\text{possible topologies}} \times \underbrace{(2N-1)}_{\text{branches per topology}} . \tag{7}$$

### 2 Extra Trees Generated in the 1D + 1L Case

We derive the number of additional gene tree topologies that are possible with one duplication and one loss (1D + 1L + Coalescence), but that cannot occur with 1D + Coalescence .

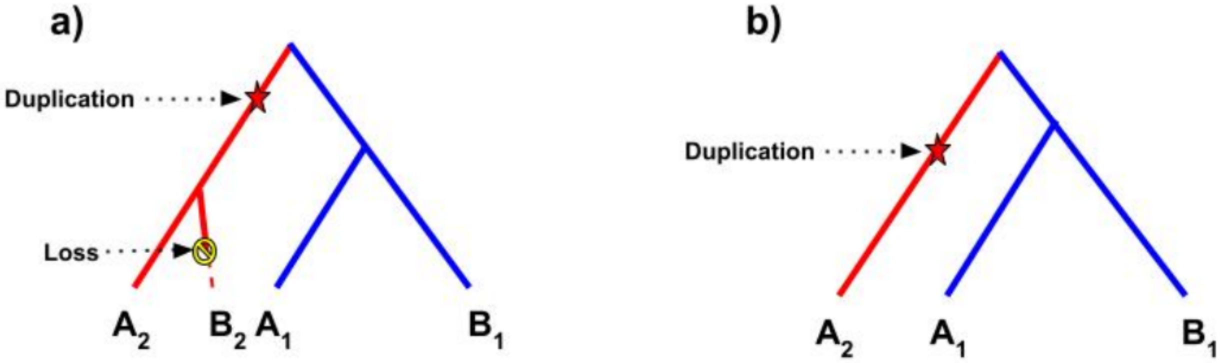


Figure 1: **Loss at the daughter locus can be explained by duplication and coalescence: a)** A duplication on the $(A_2, B_2)$ lineage and loss of lineage $B_2$ at the daughter locus, making the history of this gene tree one duplication, one loss, and coalescence. **b)** The same tree topology can also be produced by just one duplication and coalescence. The duplication of just lineage $A_2$ in this case can produce the same tree topology as in panel a.

#### 2.1 Observations

A loss in the daughter copy does not produce any new gene trees that cannot already be obtained by 1D+Coalescence. As an example, let loss occur on a branch in the daughter copy on lineage $B_2$ (Supplementary Figure 1). The resulting tree can also be explained with duplication along just the $A_2$ lineage plus coalescence to the branch subtending both copies at the parent locus. Every loss at the daughter locus (except loss on the root branch) can be explained this way. Hence, only losses landing on the parent locus create genuinely new topologies. Therefore, adding a single loss after a duplication will only change the attachment branches (parent locus) and will not affect the insertion branches (daughter locus).

#### 2.2 Possible size of the remaining parent lineage

On any parent locus with $N$ species there are three kind of branches, with losses landing on each of these branches producing different classes of lineages based on clade size, $k$ .

1. A loss on an external branch will give a remaining clade of size $k = N - 1$.
2. A loss on an internal branch subtending $M$ taxa will give a remaining clade of size $k = N - M$.
3. A loss on a root branch will result in a clade of size $k = 0$. We will ignore this case as it is the loss of the entire parent locus.

Thus we consider that the parent copy can produce clade of sizes $S \in \{1, ..., N - 1\}$.

#### 2.3 Extra attachment branches created by a loss

For each surviving clade in a parent locus of size $k$ there are $(2k - 3)!!$ rooted gene trees. Each tree contains $(2k - 1)$ branches excluding the root, and 1 root branch. Thus, the total number of additional possible attachment branches in the parent locus created by a single loss after a duplication is:

$$\sum_{k=1}^{N-1} \binom{N}{k} (2k-3)!!(2k-1) \tag{8}$$

This equation is counting all the possible subsets that can be formed with one loss (explained in section 2.2), along with all the trees that can be formed from those subsets and all the branches within those trees.

#### 2.4 Number of possible insertion branches

Since a loss landing on a daughter copy does not add any new trees that 1D+Coalescence cannot produce, the number of possible insertion branches remains the same (see section 2.1)

#### 2.5 Loss of the daughter copy

Extra lineages are also created when the loss lands on the root branch of the daughter locus. This creates no lineages at the daughter locus, resulting in the tree at the parent locus being the final gene tree. Thus for $N$ species we add $(2N-3)!!$ extra gene trees from parent locus alone if there is loss of the entire daughter copy.

#### 2.6 Final expression

For each attachment branch generated by loss, we must multiply it with the number of possible insertion branches to get the total number of extra trees generated by adding 1 loss:

$$\text{ExtraGeneTrees}(N) = \left( \sum_{k=1}^{N-1} \binom{N}{k} (2k-3)!!(2k-1) \right) \left( \sum_{k=1}^{N} \binom{N}{k} (2k-3)!! \right) + (2N-3)!! \tag{9}$$

##### 2.6.1 Example when $N = 2$

When $N = 2$, we obtain the following extra trees:

$$\left( \underbrace{\{A_1, B_1\}}_{\text{losses in leaves}} \right) \times \left( \underbrace{\{A_2, B_2, (A_2, B_2)\}}_{\text{all duplication lineages}} \right) + \underbrace{\emptyset}_{\text{when daughter copy is lost}} \times \underbrace{\{(A_1, B_1)\}}_{\text{all gene tree topologies in parent copy}} = 2 \times 3 + 1 = 7$$

1. **When duplication happens in root and loss happens in leaves**
   - $(A_1, (A_2, B_2))$
   - $(B_1, (A_2, B_2))$
2. **When duplication happens in $A_2$ and loss happens in leaves**
   - $(A_1, A_2)$
   - $(B_1, A_2)$
3. **When duplication happens in $B_2$ and loss happens in leaves**
   - $(A_1, B_2)$
   - $(B_1, B_2)$
4. **Loss of entire daughter copy**
   - $(A_1, B_1)$

Table 1: Number of extra trees generated by 1D+1L+Coalescence $N = 2$ to $N = 6$

| $N$ | **Extra Attachment Branches** | **Number of Possible Duplication Branches** | **Loss of Daughter Copy** | **Extra Trees** |
|---|---|---|---|---|
| 2 | 2 | 3 | 1 | 7 |
| 3 | 12 | 9 | 3 | 111 |
| 4 | 82 | 37 | 15 | 3049 |
| 5 | 710 | 225 | 105 | 159855 |
| 6 | 7596 | 1881 | 945 | 14289021 |

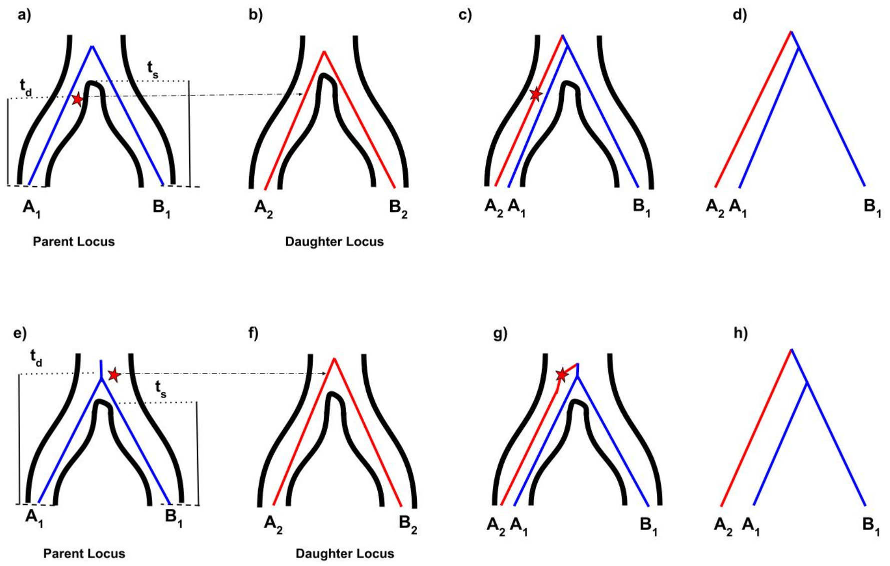


Figure 2: Ways to generate $(A_2, (A_1, B_1))$. **a-d)** When the time of duplication is less than the time of speciation ($t_d < t_s$) and the duplication inserts on the $A$ lineage, the only way to obtain the tree $(A_2, (A_1, B_1))$ is if the daughter copy $A_2$ fails to coalesce with the parent copy $A_1$ before $t_s$ and enters the ancestral population. Once $A_2$ enters the ancestral population, the daughter copy can attach with $(A_1, B_1)$ generating the required tree. **e-h)**When the time of duplication is more than the time of speciation ($t_d \geq t_s$) and the duplication occurs in the $(A, B)$ population, the only way to obtain the tree $(A_2, (A_1, B_1))$ is if the duplication inserts on daughter branch $A_2$. When the daughter copy attaches with the parent copy above the coalescence time at that locus (i.e. the branch subtending $(A_1, B_1)$), the required tree is generated.

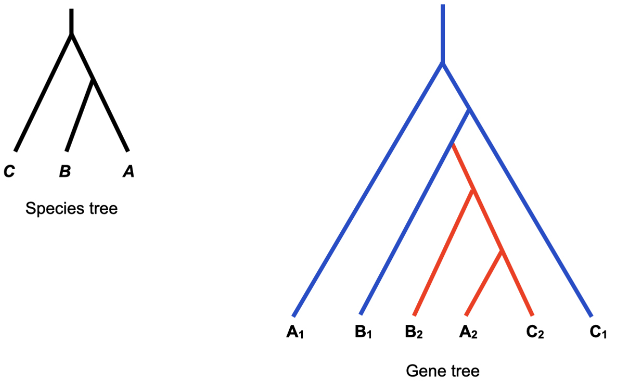


Figure 3: **One duplication and coalescence in a three-species tree.** Here, a single duplicate has occurred and attached to the branch subtending lineages $A_2, B_2, C_2$ at the daughter locus, with the gene tree at this locus being discordant with the species tree (shown to the left). The duplication has attached to the $B_1$ lineage at the parent locus, which also has a discordant gene tree. In total, standard DL reconciliation would infer 4 duplications and 8 losses given this gene tree and species tree.

### 3 Expected Internal Branch Lengths

In this section we go through the derivations for all expected internal branch lengths given in Table 3 in the main text.

#### 3.1 Tree 1: $((A_1, B_1), (B_2, A_2))$

##### 3.1.1 Problem setup

Given species tree $(A, B)$ with speciation time $t_s$ and duplication rate $\lambda$, our goal is to calculate the expected internal branch lengths of tree 1, $((A_1, B_1), (B_2, A_2))$, when this topology is generated with precisely one duplication ($n = 1$).

##### 3.1.2 Definition

Let $t_j$ and $t_i$ be the time of coalescence at parent and daughter locus, respectively, which occurs in the common population of $A$ and $B$.

$$\begin{aligned} t_j &= t_s + X_j \qquad X_j \overset{\text{iid}}{\sim} \text{Exp}(1) \\ t_i &= t_s + X_i \qquad X_i \overset{\text{iid}}{\sim} \text{Exp}(1) \end{aligned} \tag{10}$$

Let $t_k$ be the time at which the daughter lineage coalesces with the parent tree; for this topology, $t_k$ represents the coalescence time of $(A_2, B_2)$ with $(A_1, B_1)$. By definition, $t_k$ must happen above $t_s$, $t_j$, and $t_i$.

Define event $\mathcal{E}_1$ (the generation of this tree with one duplication) as:

$$\mathcal{E}_1 = \{((A_1, B_1), (A_2, B_2)) \mid \lambda, t_s, n = 1\} \tag{11}$$

There are two internal branches in this tree. We define the first branch ($b_1$) as the branch leading to the parent clade−$(A_1, B_1)$. Similarly, we define the second branch ($b_2$) as the branch leading to the parent leading to the daughter clade−$(A_2, B_2)$.

$$\begin{aligned} b_1 &= t_k - t_j \\ b_2 &= t_k - t_i \end{aligned} \tag{12}$$

(see also Figure 2 in the main text). Therefore, we want to find expected branch length of $b_1$ and $b_2$ given respectively by $\mathbb{E}[b_1|\mathcal{E}_1]$ and $\mathbb{E}[b_2|\mathcal{E}_1]$.

##### 3.1.3 Ordering of time

Given this tree and our assumptions, there is only one period in which duplication can occur. In later trees there will be more than one such period (or "regime"), but in this tree topology the time of duplication ($t_d$) must be ancestral to the time of coalescence at the daughter locus ($t_i$) and more recent than the time of coalescence at the parent locus ($t_j$). As stated earlier, the time of coalescence of parent and daughter sub-trees ($t_k$) must occur after time of coalescence at the parent locus. Thus the ordering of time of events is: $t_s < t_i < t_d < t_j < t_k$.

##### 3.1.4 Prior distribution of the gene tree branch lengths

Without conditioning on a number of duplications, there are two probability distributions we can define:

1. The density of $t_j$ and $t_i$ under the coalescent model are:

$$\begin{aligned} f_{t_j}(t_j) &= e^{-(t_j - t_s)} \\ f_{t_i}(t_i) &= e^{-(t_i - t_s)} \end{aligned} \tag{13}$$

2. The probability of duplication happening at any particular time ($t_d = d$) is given by:

$$\Pr(t_d = d) = \frac{1}{t_j} \tag{14}$$

since the duplication event has to happen at the parent locus in our model.

###### 3.1.5 Conditional

We now must define the conditions required to get this particular tree:

1. In order to get this overall tree topology, the duplicated lineage from the daughter tree must not coalesce with the parent lineage before $t_j$ (i.e. $t_j < t_k$). Given a fixed time of coalescence of the parent sub-tree, $t_j$, there are two coalescent events that must not happen in between $t_d$ and $t_j$: $(A_2, B_2)$ with either $A_1$ or $B_1$. The probability of these events not happening is given by:
$$e^{-2(t_j - t_d)} \tag{15}$$
After that, the time of coalescence of parent locus with daughter locus above $t_j$ is an exponential process with rate 1:
$$e^{-(t_k - t_j)} \tag{16}$$

2. $\mathcal{E}_1$ also has the requirement that the number of duplication events is one. This adds two more conditions. There must be precisely one duplication at the parent locus. Since we are allowing duplication to happen until height $t_j$, the total branch length in the parent sub-tree is given by $2t_j$. Thus, under a Poisson model of duplicative mutation, the probability of one duplication happening at the parent locus is given by
$$\Pr(n = 1|\lambda, t_j) = 2\lambda t_j e^{-2\lambda t_j} \tag{17}$$
Similarly, another condition is that there must be no additional duplications on the daughter locus. Since the branch length on the relevant daughter sub-tree is determined by the time of duplication $t_d$ and the duplication landing on the branch subtending the $(A_2, B_2)$ clade, this sub-tree has three branches. The first branch is the subtending branch of length $t_d - t_i$. The second and third branches are tip branches leading to $A_2$ and $B_2$ (given by $t_i$ on each side). Therefore, the total length of the daughter sub-tree at the time of duplication is $L_D = t_i + t_i + (t_d - t_i) = t_d + t_i$ Therefore, the probability of no duplication on daughter locus after there has been one duplication at the parent locus is given by
$$\Pr(n = 0|\lambda, t_d, t_i) = e^{-\lambda(t_d + t_i)} \tag{18}$$

###### 3.1.6 Posterior

The posterior of $t_k, t_d, t_j, t_i$ is proportional to the prior and conditional. We can obtain the posterior by multiplying equations 13 to 18.

$$\Pr(t_j, t_k, t_d, t_i \mid \mathcal{E}_1) \propto \underbrace{e^{-(t_j - t_s)} e^{-(t_i - t_s)} \frac{1}{t_j}}_{\text{prior}} \underbrace{\cdot e^{-2(t_j - t_d)} e^{-(t_k - t_j)} 2\lambda t_j e^{-2\lambda t_j} e^{-\lambda(t_d + t_i)}}_{\text{conditional}} \tag{19}$$

###### 3.1.7 Solving

We can now solve equation (19) to get the distribution of internal branches. For simplicity, let $\gamma = t_d - t_i$ and $\theta = t_j - t_d$; by definition, $b_1 = t_k - t_j$. Replacing terms in equation (19) we get:

$$\Pr(t_j, t_k, t_d, t_i \mid \mathcal{E}_1) \propto e^{-(t_i + \gamma + \theta - t_s)} e^{-(t_i - t_s)} \quad \cdot e^{-2(t_i + \gamma + \theta - t_d)} e^{-((t_i + \gamma + \theta + b_1) - t_i + \gamma + \theta)} e^{-2\lambda(t_i + \gamma + \theta)} e^{-\lambda(t_d + t_i)} \tag{20}$$

Now, we can solve for $\theta$, $\gamma$, and $b_1$, which lead to $b_2$:

$$\theta \sim \text{Exp}(3 + 2\lambda) \tag{21}$$

$$\gamma \sim \text{Exp}(1 + 3\lambda) \tag{22}$$

$$b_1 \sim \text{Exp}(1) \tag{23}$$

$$\boxed{\mathbb{E}[b_1|\mathcal{E}_1] = 1} \tag{24}$$

$$b_2 = \gamma + \theta + b_1 \tag{25}$$

$$\mathbb{E}[b_2] = \mathbb{E}[\gamma + \theta + b_1] = \mathbb{E}[\gamma] + \mathbb{E}[\theta] + \mathbb{E}[b_1] = \frac{1}{1+3\lambda} + \frac{1}{3+2\lambda} + 1 \tag{26}$$

$$\boxed{\mathbb{E}[b_2|\mathcal{E}_1] = \frac{1}{1+3\lambda} + \frac{1}{3+2\lambda} + 1} \tag{27}$$

Therefore the expectation of branches $b_1$ and $b_2$ are given by equation (24) and (27), respectively.

#### 3.2 Trees 2-3: $((A_1, (A_2, B_2)), B_1)$

##### 3.2.1 Problem setup

Given species tree $(A, B)$ with speciation time $t_s$ and duplication rate $\lambda$, our goal is to calculate the expected internal branch lengths of tree 3, $((A_1, (A_2, B_2)), B_1)$, when this topology is generated with precisely one duplication ($n = 1$). Note that the exact same calculations apply to tree 2, which has topology $((B_1, (A_2, B_2)), A_1)$.

##### 3.2.2 Definition

Let $t_j$ and $t_i$ be the time of coalescence at parent and daughter locus, respectively, in the common population of $A$ and $B$.

$$\begin{aligned} t_j &= t_s + X_j \qquad X_j \overset{\text{iid}}{\sim} \text{Exp}(1) \\ t_i &= t_s + X_i \qquad X_i \overset{\text{iid}}{\sim} \text{Exp}(1) \end{aligned} \tag{28}$$

Let $t_k$ be the time at which the daughter lineage coalesces with the parent tree; for this topology, $t_k$ represents the coalescence time of $(A_2, B_2)$ with $A_1$. By definition, $t_k$ must happen above $t_s$ and $t_i$ but below $t_j$

Define event $\mathcal{E}_2$ (the generation of this tree with one duplication) as:

$$\mathcal{E}_2 = \{((A_1, (A_2, B_2)), B_1) \mid \lambda, t_s, n = 1\} \tag{29}$$

There are two internal branches in this tree. We define the first branch ($b_{lower}$) as the branch leading to the daughter clade−$(A_2, B_2)$. Similarly, we define the second branch ($b_{upper}$) as the branch leading to the clade−$(A_1, (A_2, B_2))$.

$$\begin{aligned} b_{lower} &= t_k - t_i \\ b_{upper} &= t_j - t_k \end{aligned} \tag{30}$$

(see also Figure 2 in the main text). Therefore, we want to find expected branch length of $b_{lower}$ and $b_{upper}$ given respectively by $\mathbb{E}[b_{lower}|\mathcal{E}_2]$ and $\mathbb{E}[b_{upper}|\mathcal{E}_2]$.

##### 3.2.3 Ordering of time

There is only one regime for this tree. In this regime the time of duplication($t_d$) must be ancestral to the time of coalescence at the daughter locus ($t_i$) and more recent than the time of coalescence at the parent locus ($t_j$). The time of coalescence of parent and daughter sub-trees ($t_k$) must occur before time of coalescence at the parent locus. Thus the ordering of time is: $t_s < t_i < t_d < t_k < t_j$.

##### 3.2.4 Prior distribution of the gene tree branch lengths

Without conditioning on a number of duplications, there are two probability distributions we can define:

1. The density of $t_j$ and $t_i$ under the coalescent model are:

$$\begin{aligned} f_{t_j}(t_j) &= e^{-(t_j - t_s)} \\ f_{t_i}(t_i) &= e^{-(t_i - t_s)} \end{aligned} \tag{31}$$

2. The probability of duplication happening at any particular time ($t_d = d$) is given by:

$$\Pr(t_d = d) = \frac{1}{t_j} \tag{32}$$

##### 3.2.5 Conditional

1. In order to get this overall tree topology, the duplicated lineage from the daughter tree must coalesce with parent lineage before $t_j$ (i.e. $t_j > t_k$). Given a fixed time of coalescence of the parent sub-tree, $t_j$, there are two coalescent events that must happen in between $t_d$ and $t_j$: $(A_2, B_2)$ with either $A_1$ or $B_1$. The probability of these events happening is given by:
$$e^{-2(t_k - t_d)} \tag{33}$$
2. $\mathcal{E}_2$ also has the condition that the number of duplication events is one. This adds two more conditions.
There must be precisely one duplication at the parent locus. Since we are allowing duplication to happen until height $t_j$, the total branch length in the parent sub-tree is given by $2t_j$. Thus, the probability of one duplication landing on the parent locus is given by
$$\Pr(n = 1|\lambda, t_j) = 2\lambda t_j e^{-2\lambda t_j} \tag{34}$$
Similarly, another condition is that there must be no additional duplication on the daughter locus after the first duplication. Since the branch length on the relevant daughter sub-tree is determined by the time of duplication $t_d$ and the duplication landing on the branch subtending the $(A_2, B_2)$ clade, this sub-tree has three branches. The first branch is the subtending branch of length $t_d - t_i$. The second and third branches are tip branches leading to $A_2$ and $B_2$ (given by $t_i$ on each side). Therefore, the total length of the daughter sub-tree at the time of duplication is $L_D = t_i + t_i + (t_d - t_i) = t_d + t_i$ Therefore, the probability of no duplication on daughter locus after there has been one duplication at the parent locus is given by
$$\Pr(n = 0|\lambda, t_d, t_i) = e^{-\lambda(t_d + t_i)} \tag{35}$$

##### 3.2.6 Posterior

The posterior of $t_k, t_d, t_j, t_i$ is proportional to the prior and conditional. We can obtain the posterior by multiplying equations 32 to 36.

$$\Pr(t_j, t_k, t_d, t_i \mid \mathcal{E}_2) \propto \underbrace{e^{-(t_j - t_s)} e^{-(t_i - t_s)} \frac{1}{t_j}}_{\text{prior}} \cdot \underbrace{e^{-2(t_k - t_d)} 2\lambda t_j e^{-2\lambda t_j} e^{-\lambda(t_d + t_i)}}_{\text{conditional}} \tag{36}$$

##### 3.2.7 Solving

We can now solve equation (36) to get distribution of internal branches. For simplicity, let $\gamma = t_d - t_i$ , $\theta = t_i - t_s$ ,and $\alpha = t_k - t_d$; by definition, $b_{upper} = t_j - t_k$. Replacing terms in equation (36) and rearranging we get:

$$\Pr(t_j, t_k, t_d, t_i \mid \mathcal{E}_2) \propto e^{-(2+4\lambda)(t_i - t_s)} e^{-(1+3\lambda)(t_d - t_i)} e^{-(3+2\lambda)(t_k - t_d)} e^{-(1+2\lambda)(t_j - t_k)} \tag{37}$$

Now, we can solve for: $\gamma, \theta, \alpha$ are $b_1$, which lead to $b_{lower}$ and $b_{upper}$:

$$\theta \sim \text{Exp}(2 + 4\lambda) \tag{38}$$
$$\gamma \sim \text{Exp}(1 + 3\lambda) \tag{39}$$
$$\alpha \sim \text{Exp}(3 + 2\lambda) \tag{40}$$
$$b_{upper} \sim \text{Exp}(1 + 2\lambda) \tag{41}$$

$$\boxed{\mathbb{E}[b_{upper}|\mathcal{E}_2] = \frac{1}{1 + 2\lambda}} \tag{42}$$

$$\mathbb{E}[b_{lower}] = \mathbb{E}[\gamma + \alpha] = \mathbb{E}[\gamma] + \mathbb{E}[\alpha] = \frac{1}{1 + 3\lambda} + \frac{1}{3 + 2\lambda} \tag{43}$$

$$\boxed{\mathbb{E}[b_{lower}|\mathcal{E}_2] = \frac{1}{1 + 3\lambda} + \frac{1}{3 + 2\lambda}} \tag{44}$$

Therefore the expectation of branches $b_{lower}$ and $b_{upper}$ is given by equation (44) and (42) respectively .

#### 3.3 Trees 4-5: $((B_1, B_2), A_1)$

##### 3.3.1 Problem setup

Given species tree $(A, B)$ with speciation time $t_s$ and duplication rate $\lambda$, our goal is to calculate the expected internal branch lengths of tree 5, $((B_1, B_2), A_1)$, when this topology is generated with precisely one duplication ($n = 1$). Note that the exact same calculations apply to tree 4, which has topology $((A_1, A_2), B_1)$.

##### 3.3.2 Definition

Let $t_j$ be the time of coalescence at the parent locus (i.e. $A_1$ with $B_1$), in the common population of $A$ and $B$.

$$t_j = t_s + X_j \qquad X_j \overset{\text{iid}}{\sim} \text{Exp}(1) \tag{45}$$

Let $t_k$ be the time at which the daughter lineage coalesces with the parent tree; for this topology, $t_k$ represents the coalescence time of $B_2$ with $B_1$.

Define event $\mathcal{E}_4$ (the generation of this tree with one duplication) as:

$$\mathcal{E}_4 = \{((B_1, B_2), A_1) \mid \lambda, t_s, n = 1\} \tag{46}$$

Based on the definition of $t_j$ and $t_k$ the internal branch is defined as:

$$b = t_j - t_k \tag{47}$$

Therefore, we want to find $\mathbb{E}[b \mid \mathcal{E}_4]$.

##### 3.3.3 Ordering of time

There are three regimes for this tree based on the relative timing of duplication and speciation:

**Regime 1 ($\mathcal{R}_1$) :** The time of duplication ($t_d$) is more recent than the time of speciation ($t_s$), and the time of coalescence of the parent and daughter sub-trees ($t_k$) occurs before the time of speciation. Thus, the ordering of time is $t_d < t_k < t_s < t_j$.

**Regime 2 ($\mathcal{R}_2$) :** The time of duplication ($t_d$) is more recent than the time of speciation ($t_s$), and the time of coalescence of the parent and daughter sub-trees ($t_k$) occurs after the time of speciation, but before $t_j$. Thus, the ordering of time is $t_d < t_s < t_k < t_j$.

**Regime 3 ($\mathcal{R}_2$) :** The time of duplication ($t_d$) occurs after the time of speciation ($t_s$), and the time of coalescence of the parent and daughter sub-trees ($t_k$) occurs after the time of speciation, but before $t_j$. Thus, the ordering of time is $t_s < t_d < t_k < t_j$

##### 3.3.4 Prior distribution of the gene tree branch lengths

Without conditioning on a number of duplications, there are two probability distributions we can define:

1. The density of $t_j$ under the coalescent model is:

$$f_{t_j}(t_j) = e^{-(t_j - t_s)} \tag{48}$$

2. The probability of duplication happening at any particular time ($t_d = d$) is given by:

$$\Pr(t_d = d) = \frac{1}{t_j} \tag{49}$$

##### 3.3.5 Conditional

1. In **regime 1** ($\mathcal{R}_1$), to get this tree topology, the duplicated lineage from the daughter tree coalesces with the parent lineage before speciation. Thus, the time of coalescence of $B_2$ with $B_1$ (given by $t_k$) is an exponential process with rate 1 starting at time $t_d$, and is given by:

$$e^{-(t_k - t_d)} \tag{50}$$

2. In **regime 2** ($\mathcal{R}_2$), to get this tree topology, the duplicated lineage from the daughter tree must not coalesce before speciation, instead coalescing with $B_1$ in the common population of $A$ and $B$. In the common population, the time of coalescence of $A_1$ and $B_1$ (given by $t_j$) is already fixed. Therefore, there are two possible pairs, $(A_1, B_2)$ and $(B_1, B_2)$, that can coalesce between $t_s$ and $t_j$, which makes the rate of the exponential process equal to 2. Hence, the density of $t_k$ in regime 2 is given by:

$$e^{-(t_s - t_d)} e^{-2(t_k - t_s)} \tag{51}$$

3. In **regime 3** ($\mathcal{R}_3$), to get this tree topology, the duplication occurs after speciation (in the common population of $A$ and $B$), and must land on branch $B_2$ at the daughter locus. This means that by the time of duplication, the daughter lineages must not have coalesced. After that there is then an exponential process between $t_d$ and $t_k$ with rate 2 (similar to regime 2). Hence, the density of $t_k$ in regime 3 is given by:

$$e^{-(t_d - t_s)} e^{-2(t_k - t_d)} \tag{52}$$

4. $\mathcal{E}_4$ also has the requirement that the number of duplication events is one. This adds two more conditions. There must be precisely one duplication at the parent locus. Since we are allowing duplication to happen until height $t_j$, the total branch length in the parent sub-tree is given by $2t_j$. Thus, the probability of one duplication landing on the parent locus is given by:

$$\Pr(n = 1 | \lambda, t_j) = 2\lambda t_j e^{-2\lambda t_j} \tag{53}$$

Similarly, another condition is that there must be no additional duplication on the daughter locus. Since the branch length on the relevant daughter sub-tree is determined by the time of duplication $t_d$ and the duplication landing on the branch subtending the $B_2$ clade, this sub-tree has one branch. Therefore, the probability of no duplication on the daughter locus after there has been one duplication at the parent locus is given by:

$$\Pr(n = 0 | \lambda, t_d) = e^{-\lambda t_d} \tag{54}$$

##### 3.3.6 Posterior

The posterior of $t_k, t_d, t_j$ for each of the regimes is proportional to the prior and conditional of that regime.
For **regime 1** the posterior is proportional to the following:

$$\Pr(t_j, t_k, t_d \mid \mathcal{E}_4, \mathcal{R}_1) \propto \underbrace{e^{-(t_j - t_s)} \frac{1}{t_j}}_{\text{prior}} \cdot \underbrace{2\lambda t_j e^{-2\lambda t_j} e^{-\lambda t_d} e^{-(t_k - t_d)}}_{\text{conditional}} \tag{55}$$

For **regime 2** the posterior is proportional to the following:

$$\Pr(t_j, t_k, t_d \mid \mathcal{E}_4, \mathcal{R}_2) \propto \underbrace{e^{-(t_j - t_s)} \frac{1}{t_j}}_{\text{prior}} \cdot \underbrace{2\lambda t_j e^{-2\lambda t_j} e^{-\lambda t_d} e^{-(t_s - t_d)} e^{-2(t_k - t_s)}}_{\text{conditional}} \tag{56}$$

And for **regime 3** the posterior is proportional to the following:

$$\Pr(t_j, t_k, t_d \mid \mathcal{E}_4, \mathcal{R}_3) \propto \underbrace{e^{-(t_j - t_s)} \frac{1}{t_j}}_{\text{prior}} \cdot \underbrace{2\lambda t_j e^{-2\lambda t_j} e^{-\lambda t_d} e^{-(t_d - t_s)} e^{-2(t_k - t_d)}}_{\text{conditional}} \tag{57}$$

##### 3.3.7 Solving

In this section, we first derive the expected internal branch length under each regime. We then compute the posterior weight of each regime (i.e. the probability of seeing this tree under a regime). Finally, we calculate the expected internal branch length for this tree as the sum of the expected branch lengths from each regime, each scaled by the normalized posterior weight of its corresponding regime.

**Regime 1** We can replace $t_j = b + t_k$ in equation (55)

$$\Pr(t_j, t_k, t_d \mid \mathcal{E}_4, \mathcal{R}_1) \propto e^{-(b+t_k-t_s)}\frac{1}{t_j} \cdot 2\lambda t_j e^{-2\lambda(b+t_k)} e^{-\lambda t_d} e^{-(t_k-t_d)} \tag{58}$$

Rearranging expressions that depends upon $b$ we get:

$$\Pr(b \mid \mathcal{E}_4, \mathcal{R}_1) \cdot \Pr(t_k, t_d \mid \mathcal{E}_4, \mathcal{R}_1) = \int_{t_s-t_k}^{\infty} e^{-(1+2\lambda)b} db \cdot \int_0^{t_s} e^{-2(1+\lambda)t_k} dt_k \cdot \int_0^{t_k} e^{(1-\lambda)t_d} dt_d \tag{59}$$

Now solving equation (59) we get:

$$\begin{aligned}
\Pr(b \mid \mathcal{E}_4, \mathcal{R}_1) \cdot \Pr(t_k, t_d \mid \mathcal{E}_4, \mathcal{R}_1) &= \int_0^{t_s} \frac{e^{-(1+2\lambda)(t_s-t_k)}}{1+2\lambda} \cdot e^{-2(1+\lambda)t_k} dt_k \cdot \int_0^{t_k} e^{(1-\lambda)t_d} dt_d \\
&= \int_0^{t_s} \frac{e^{-(1+2\lambda)(t_s-t_k)}}{1+2\lambda} \cdot e^{-2(1+\lambda)t_k} \cdot \frac{e^{(1-\lambda)t_k} - 1}{1-\lambda} dt_k \\
&= \frac{e^{-(1+2\lambda)t_s}}{(1+2\lambda)(1-\lambda)} \int_0^{t_s} \left(e^{-\lambda t_k} - e^{-t_k}\right) dt_k
\end{aligned}$$

Hence,

$$\mathbb{E}[t_k \mid \mathcal{R}_1] = \frac{\int_0^{t_s} t_k \left(e^{-\lambda t_k} - e^{-t_k}\right) dt_k}{\int_0^{t_s} \left(e^{-\lambda t_k} - e^{-t_k}\right) dt_k}$$

Therefore,

$$\boxed{\mathbb{E}[b \mid \mathcal{R}_1] = \frac{\int_{t_s-t_k}^{\infty} b e^{-(1+2\lambda)b}\, db}{\int_{t_s-t_k}^{\infty} e^{-(1+2\lambda)b}\, db} = t_s - \mathbb{E}[t_k] + \frac{1}{1+2\lambda} = t_s - \frac{\int_0^{t_s} t_k \left(e^{-\lambda t_k} - e^{-t_k}\right) dt_k}{\int_0^{t_s} \left(e^{-\lambda t_k} - e^{-t_k}\right) dt_k} + \frac{1}{1+2\lambda}} \tag{60}$$

The probability of seeing this tree under the conditions of $\mathcal{R}_1$ can also be derived directly from equation (58):

$$P_1 \propto e^{t_s} \int_{t_s-t_k}^{\infty} e^{-(1+2\lambda)b} db \cdot \int_0^{t_s} e^{-2(1+\lambda)t_k} dt_k \cdot \int_0^{t_k} e^{(1-\lambda)t_d} dt_d \tag{61}$$

**Regime 2** We can replace $t_j = b + t_k$ in equation (56)

$$\Pr(t_j, t_k, t_d \mid \mathcal{E}_4, \mathcal{R}_2) \propto e^{-(b+t_k-t_s)}\frac{1}{t_j} \cdot 2\lambda t_j e^{-2\lambda(b+t_k)} e^{-\lambda t_d} e^{-(t_s-t_d)} e^{-2(t_k-t_s)} \tag{62}$$

Rearranging expressions that depend upon $b$ we get:

$$\Pr(b \mid \mathcal{E}_4, \mathcal{R}_2) \cdot \Pr(t_k, t_d \mid \mathcal{E}_4, \mathcal{R}_2) = \int_0^{\infty} e^{-(1+2\lambda)b} db \cdot \int_{t_s}^{\infty} e^{-(3+2\lambda)t_k} dt_k \cdot \int_0^{t_s} e^{(1-\lambda)t_d} dt_d \tag{63}$$

Because in equation (63), $b$ is independent of other terms:

$$\boxed{\mathbb{E}[b \mid \mathcal{R}_2] = \frac{\displaystyle\int_0^{\infty} be^{-(1+2\lambda)b}\,db}{\displaystyle\int_0^{\infty} e^{-(1+2\lambda)b}\,db} = \frac{1}{1+2\lambda}} \tag{64}$$

The probability of seeing this tree under conditions of $\mathcal{R}_2$ can be derived from equation (62):

$$P_2 \propto e^{2t_s} \int_0^{\infty} e^{-(1+2\lambda)b} db \cdot \int_{t_s}^{\infty} e^{-(3+2\lambda)t_k} dt_k \cdot \int_0^{t_s} e^{(1-\lambda)t_d} dt_d \tag{65}$$

**Regime 3** We can replace $t_j = b + t_k$ in equation (57)

$$\Pr(t_j, t_k, t_d \mid \mathcal{E}_4, \mathcal{R}_3) \propto e^{-(b+t_k-t_s)} \frac{1}{t_j} \cdot 2\lambda t_j e^{-2\lambda(b+t_k)} e^{-\lambda t_d} e^{-(t_d-t_s)} e^{-2(t_k-t_d)} \tag{66}$$

Rearranging expressions that depend upon $b$ we get:

$$\Pr(b \mid \mathcal{E}_4, \mathcal{R}_3) \cdot \Pr(t_k, t_d \mid \mathcal{E}_4, \mathcal{R}_3) = \int_0^{\infty} e^{-(1+2\lambda)b} db \cdot \int_{t_s}^{\infty} e^{-(3+2\lambda)t_k} dt_k \cdot \int_{t_s}^{t_k} e^{(1-\lambda)t_d} dt_d \tag{67}$$

Because in equation (67) , $b$ is independent of other terms:

$$\boxed{\mathbb{E}[b \mid \mathcal{R}_3] = \frac{\displaystyle\int_0^{\infty} be^{-(1+2\lambda)b}\,db}{\displaystyle\int_0^{\infty} e^{-(1+2\lambda)b}\,db} = \frac{1}{1+2\lambda}} \tag{68}$$

The probability of seeing this tree under conditions of $\mathcal{R}_2$ can be derived from equation (67):

$$P_3 \propto e^{2t_s} \int_0^{\infty} e^{-(1+2\lambda)b} db \cdot \int_{t_s}^{\infty} e^{-(3+2\lambda)t_k} dt_k \cdot \int_{t_s}^{t_k} e^{(1-\lambda)t_d} dt_d \tag{69}$$

###### 3.3.7.1 Weighted solution

Now that we have the expected branch length for each regime (given by (60), (64), and (68)) and the corresponding posterior weights of the regimes (given by (61), (65), and (69)), we can obtain $\mathbb{E}[b \mid \mathcal{E}_4]$ by multiplying the expected branch length for each regime by the corresponding posterior weight of that regime. However, since $P_1$, $P_2$, and $P_3$ do not sum to 1 (they do not have to), we need to normalize them to ensure proper scaling. We scale by summing all three and dividing by this sum, giving us normalized posterior weights, $\tilde{P}_1$, $\tilde{P}_2$, and $\tilde{P}_3$.

We can now give the final expected internal branch length for this tree:

$$\boxed{\mathbb{E}[b \mid \mathcal{E}_4] = \tilde{P}_1 \cdot \mathbb{E}[b \mid \mathcal{R}_1] + \tilde{P}_2 \cdot \mathbb{E}[b \mid \mathcal{R}_2] + \tilde{P}_3 \cdot \mathbb{E}[b \mid \mathcal{R}_3]} \tag{70}$$

#### 3.4 Trees 6-7: $((A_1, B_1), B_2)$

##### 3.4.1 Problem setup

Given species tree $(A, B)$ with speciation time $t_s$ and duplication rate $\lambda$, our goal is to calculate the expected internal branch lengths of tree 7, $((A_1, B_1), B_2)$, when this topology is generated with precisely one duplication ($n = 1$). Note that the exact same calculations apply to tree 6, which has topology $((A_1, B_1), A_2)$.

##### 3.4.2 Definition

Let $t_j$ be the time of coalescence at the parent locus (i.e. $A_1$ with $B_1$), in the common population of $A$ and $B$.

$$t_j = t_s + X_j \qquad X_j \overset{\text{iid}}{\sim} \text{Exp}(1) \tag{71}$$

Let $t_k$ be the time at which the daughter lineage coalesces with the parent tree; for this topology, $t_k$ represents the coalescence time of $(A_1, B_1)$ with $A_2$.

Define event $\mathcal{E}_6$ (the generation of this tree with one duplication) as:

$$\mathcal{E}_6 = \{((A_1, B_1), B_2) \mid \lambda, t_s, n = 1\} \tag{72}$$

Based on the definition of $t_j$ and $t_k$ the internal branch is defined as:

$$b = t_k - t_j \tag{73}$$

Therefore, we want to find $\mathbb{E}[b \mid \mathcal{E}_6]$.

##### 3.4.3 Ordering of time

Given this tree and our assumptions, there are two regimes in which duplication can occur. These regimes are based on the relative timing of duplication and speciation.

**Regime 1 ($\mathcal{R}_1$) :** The time of duplication ($t_d$) is more recent than the time of speciation ($t_s$), and the time of coalescence of the parent and daughter sub-trees ($t_k$) occurs after the time of speciation and $t_j$. Thus, the ordering of time is $t_d < t_s < t_j < t_k$.

**Regime 2 ($\mathcal{R}_2$) :** The time of duplication ($t_d$) is ancestral than the time of speciation ($t_s$) but is more recent compared to $t_j$. And the time of coalescence of the parent and daughter sub-trees ($t_k$) occurs after the time $t_j$. Thus, the ordering of time is $t_s < t_d < t_j < t_k$.

##### 3.4.4 Prior distribution of the gene tree branch lengths

Without conditioning on a number of duplications, there are two probability distributions we can define:

1. The density of $t_j$ under the coalescent model is:

$$f_{t_j}(t_j) = e^{-(t_j - t_s)} \tag{74}$$

2. The probability of duplication happening at any particular time ($t_d = d$) is given by:

$$\Pr(t_d = d) = \frac{1}{t_j} \tag{75}$$

##### 3.4.5 Conditional

1. In **regime 1 ($\mathcal{R}_1$)**, to get this tree topology, the duplicated lineage from the daughter tree must not coalesce before speciation, instead coalescing with $(A_1, B_1)$ in the common population of $A$ and $B$. In the common population, the time of coalescence of $A_1$ and $B_1$ (given by $t_j$) is already fixed. Therefore, there are two possible pairs, $(A_1, B_2)$ and $(B_1, B_2)$, that must not coalesce between $t_s$ and $t_j$, which makes the rate of the exponential process equal to 2. Then, the time of coalescence of the parent and daughter (given by $t_k$) follows an exponential process with rate 1 starting from $t_j$. Hence, the density of $t_k$ in regime 1 is given by:

$$e^{-(t_s - t_d)} e^{-2(t_j - t_s)} e^{-(t_k - t_j)} \tag{76}$$

2. In **regime 2 ($\mathcal{R}_2$)**, to get this tree topology, the duplication occurs after speciation (in the common population of $A$ and $B$), and must land on branch $B_2$ at the daughter locus. This means that by the time of duplication, the daughter lineages must not have coalesced. After that, there is an exponential process between $t_d$ and $t_j$ with

rate 2 (similar to regime 1). Then, the time of coalescence of the parent and daughter (given by $t_k$) follows an exponential process with rate 1 starting from $t_j$. Hence, the density of $t_k$ in regime 2 is given by:

$$e^{-(t_d-t_s)}e^{-2(t_j-t_d)}e^{-(t_k-t_j)} \tag{77}$$

3. $\mathcal{E}_6$ also has the requirement that the number of duplication events is one. This adds two more conditions. There must be precisely one duplication at the parent locus. Since we are allowing duplication to happen until height $t_j$, the total branch length in the parent sub-tree is given by $2t_j$. Thus, the probability of one duplication landing on the parent locus is given by:

$$\Pr(n = 1|\lambda, t_j) = 2\lambda t_j e^{-2\lambda t_j} \tag{78}$$

Similarly, another condition is that there must be no additional duplication on the daughter locus. Since the branch length on the relevant daughter sub-tree is determined by the time of duplication $t_d$ and the duplication landing on the branch subtending the $B_2$ clade, this sub-tree has one branch. Therefore, the probability of no duplication on the daughter locus after there has been one duplication at the parent locus is given by:

$$\Pr(n = 0|\lambda, t_d) = e^{-\lambda t_d} \tag{79}$$

###### 3.4.6 Posterior

The posterior of $t_k, t_d, t_j$ for each of the regimes is proportional to the prior and conditional of that regime.
For **regime 1** the posterior is proportional to the following:

$$\Pr(t_j, t_k, t_d \mid \mathcal{E}_6, \mathcal{R}_1) \propto \underbrace{e^{-(t_j-t_s)}\frac{1}{t_j}}_{\text{prior}} \cdot \underbrace{2\lambda t_j e^{-2\lambda t_j} e^{-\lambda t_d} e^{-(t_s-t_d)} e^{-2(t_j-t_s)} e^{-(t_k-t_j)}}_{\text{conditional}} \tag{80}$$

For **regime 2** the posterior is proportional to the following:

$$\Pr(t_j, t_k, t_d \mid \mathcal{E}_6, \mathcal{R}_2) \propto \underbrace{e^{-(t_j-t_s)}\frac{1}{t_j}}_{\text{prior}} \cdot \underbrace{2\lambda t_j e^{-2\lambda t_j} e^{-\lambda t_d} e^{-(t_d-t_s)} e^{-2(t_j-t_d)} e^{-(t_k-t_j)}}_{\text{conditional}} \tag{81}$$

###### 3.4.7 Solving

In this section, we first derive the expected internal branch length under each regime. We find that the expected branch length is same for both regimes. Therefore, we use this value as the expected value of $b$.

**Regime 1** From equation (80):

$$\Pr(t_j, t_k, t_d \mid \mathcal{E}_6, \mathcal{R}_1) \propto e^{-(t_j-t_s)} \cdot 2\lambda e^{-2\lambda t_j} e^{-\lambda t_d} e^{-(t_s-t_d)} e^{-2(t_j-t_s)} e^{-(t_k-t_j)}. \tag{82}$$

$$\Pr(b \mid \mathcal{R}_1) \cdot \Pr(t_k, t_d \mid \mathcal{R}_1) \propto e^{-b} \cdot \Pr(t_k, t_d \mid \mathcal{R}_1)$$

$$\Pr(b \mid \mathcal{R}_1) \propto e^{-b}$$

**Regime 2** From equation (81):

$$\Pr(t_j, t_k, t_d \mid \mathcal{E}_6, \mathcal{R}_2) \propto e^{-(t_j-t_s)} \cdot 2\lambda e^{-2\lambda t_j} e^{-\lambda t_d} e^{-(t_d-t_s)} e^{-2(t_j-t_d)} e^{-(t_k-t_j)}. \tag{83}$$

$$\Pr(b \mid \mathcal{R}_2) \cdot \Pr(t_k, t_d \mid \mathcal{R}_2) \propto e^{-b} \cdot \Pr(t_k, t_d \mid \mathcal{R}_2)$$

$$\Pr(b \mid \mathcal{R}_2) \propto e^{-b}$$

Since $\mathbb{E}[b|\mathcal{R}_1] = \mathbb{E}[b|\mathcal{R}_2]$, the solution is the same in both regimes, and no weights are needed. Therefore:

$$\boxed{\mathbb{E}[b|\mathcal{E}_6] = 1} \tag{84}$$

#### 3.5 Trees 8-9: $((A_1, B_2), B_1)$

##### 3.5.1 Problem setup

Given species tree $(A, B)$ with speciation time $t_s$ and duplication rate $\lambda$, our goal is to calculate the expected internal branch lengths of tree 9, $((A_1, B_2), B_1)$, when this topology is generated with precisely one duplication ($n = 1$). Note that the exact same calculations apply to tree 8, which has topology $((B_1, A_2), A_1)$.

##### 3.5.2 Definition

Let $t_j$ be the time of coalescence at the parent locus (i.e. $A_1$ with $B_1$), in the common population of $A$ and $B$.

$$t_j = t_s + X_j \qquad X_j \overset{\text{iid}}{\sim} \text{Exp}(1) \tag{85}$$

Let $t_k$ be the time at which the daughter lineage coalesces with the parent tree; for this topology, $t_k$ represents the coalescence time of $A_1$ with $B_2$.

Define event $\mathcal{E}_8$ (the generation of this tree with one duplication) as:

$$\mathcal{E}_8 = \{((A_1, B_2), B_1) \mid \lambda, t_s, n = 1\} \tag{86}$$

Based on the definition of $t_j$ and $t_k$ the internal branch is defined as:

$$b = t_j - t_k \tag{87}$$

Therefore, we want to find $\mathbb{E}[b \mid \mathcal{E}_8]$.

##### 3.5.3 Ordering of time

Given this tree and our assumptions, there are two regimes in which duplication can occur. These regimes are based on the relative timing of duplication and speciation.

**Regime 1 ($\mathcal{R}_1$) :** The time of duplication ($t_d$) is more recent than the time of speciation ($t_s$), and the time of coalescence of the parent and daughter sub-trees ($t_k$) occurs after the time of speciation and but before $t_j$. Thus, the ordering of time is $t_d < t_s < t_k < t_j$.

**Regime 2 ($\mathcal{R}_2$) :** The time of duplication ($t_d$) is ancestral than the time of speciation ($t_s$) but is more recent compared to $t_j$. And the time of coalescence of the parent and daughter sub-trees ($t_k$) occurs before the time $t_j$. Thus, the ordering of time is $t_s < t_d < t_k < t_j$.

##### 3.5.4 Prior distribution of the gene tree branch lengths

Without conditioning on a number of duplications, there are two probability distributions we can define:

1. The density of $t_j$ under the coalescent model is:

$$f_{t_j}(t_j) = e^{-(t_j - t_s)} \tag{88}$$

2. The probability of duplication happening at any particular time ($t_d = d$) is given by:

$$\Pr(t_d = d) = \frac{1}{t_j} \tag{89}$$

##### 3.5.5 Conditional

1. In **regime 1 ($\mathcal{R}_1$)**, to get this tree topology, the duplicated lineage from the daughter tree must not coalesce before speciation, instead coalescing with $A_1$ in the common population of $A$ and $B$. In the common population, the time of coalescence of $A_1$ and $B_1$ (given by $t_j$) is already fixed. Therefore, there are two possible pairs, $(A_1, B_2)$ and $(B_1, B_2)$, that can coalesce between $t_s$ and $t_j$, which makes the rate of the exponential process equal to 2. Hence, the density of $t_k$ in regime 1 is given by:

$$e^{-(t_s - t_d)} e^{-2(t_k - t_s)} \tag{90}$$

2. In **regime 2 ($\mathcal{R}_2$)**, to get this tree topology, the duplication occurs after speciation (in the common population of $A$ and $B$), and must land on branch $B_2$ at the daughter locus. This means that by the time of duplication, the daughter lineages must not have coalesced. After that there is then an exponential process between $t_d$ and $t_k$ with rate 2 (similar to regime 1). Hence, the density of $t_k$ in regime 3 is given by:

$$e^{-(t_d-t_s)}e^{-2(t_k-t_d)} \tag{91}$$

3. $\mathcal{E}_8$ also has the requirement that the number of duplication events is one. This adds two more conditions. There must be precisely one duplication at the parent locus. Since we are allowing duplication to happen until height $t_j$, the total branch length in the parent sub-tree is given by $2t_j$. Thus, the probability of one duplication landing on the parent locus is given by:

$$\Pr(n=1|\lambda,t_j) = 2\lambda t_j e^{-2\lambda t_j} \tag{92}$$

Similarly, another condition is that there must be no additional duplication on the daughter locus. Since the branch length on the relevant daughter sub-tree is determined by the time of duplication $t_d$ and the duplication landing on the branch subtending the $B_2$ clade, this sub-tree has one branch. Therefore, the probability of no duplication on the daughter locus after there has been one duplication at the parent locus is given by:

$$\Pr(n=0|\lambda,t_d) = e^{-\lambda t_d} \tag{93}$$

##### 3.5.6 Posterior

The posterior of $t_k, t_d, t_j$ for each of the regime(1 and 2) is proportional to the prior and conditional of that regime.

For **regime 1** the posterior is proportional to the following term.

$$\Pr(t_j,t_k,t_d \mid \mathcal{E}_8) \propto \underbrace{e^{-(t_j-t_s)}\frac{1}{t_j}}_{\text{prior}} \cdot \underbrace{2\lambda t_j e^{-2\lambda t_j} e^{-\lambda t_d} e^{-2(t_k-t_s)} e^{-(t_s-t_d)}}_{\text{conditional}} \tag{94}$$

Similarly for **regime 2** the posterior is proportional to the following term.

$$\Pr(t_j,t_k,t_d \mid \mathcal{E}_8) \propto \underbrace{e^{-(t_j-t_s)}\frac{1}{t_j}}_{\text{prior}} \cdot \underbrace{2\lambda t_j e^{-2\lambda t_j} e^{-\lambda t_d} e^{-2(t_k-t_d)} e^{-(t_d-t_s)}}_{\text{conditional}} \tag{95}$$

##### 3.5.7 Solving

In this section, we first derive the expected internal branch length under each regime. We find that the expected branch length is same for both regimes. Therefore, we use this value as the expected value of $b$.

**Regime 1** We can replace $t_j = b + t_k$ in equation (94)

$$\Pr(t_j,t_k,t_d \mid \mathcal{E}_8,\mathcal{R}_1) \propto e^{-(t_k+b-t_s)}2\lambda e^{-2\lambda(t_k+b)}e^{-\lambda t_d}e^{-2(t_k-t_s)}e^{-(t_s-t_d)}. \tag{96}$$

Now collecting all the terms that are dependent on $b$, we get:

$$\Pr(b \mid \mathcal{R}_1)\cdot\Pr(t_k,t_d \mid \mathcal{R}_1) \propto e^{-(1+2\lambda)b}\cdot\Pr(t_k,t_d \mid \mathcal{R}_1)$$

$$\Pr(b \mid \mathcal{R}_1) \propto e^{-(1+2\lambda)b}$$

Therefore,

$$\boxed{\mathbb{E}[b \mid \mathcal{R}_1] = \frac{\displaystyle\int_0^\infty be^{-(1+2\lambda)b}\,db}{\displaystyle\int_0^\infty e^{-(1+2\lambda)b}\,db} = \frac{1}{1+2\lambda}} \tag{97}$$

**Regime 2** We can replace $t_j = b + t_k$ in equation (95)

$$\Pr(t_j, t_k, t_d \mid \mathcal{E}_8, \mathcal{R}_2) \propto e^{-(b+t_k-t_s)} \cdot 2\lambda e^{-2\lambda(b+t_k)} e^{-\lambda t_d} e^{-2(t_k-t_d)} e^{-(t_d-t_s)} \tag{98}$$

Now collecting all the terms that are dependent on $b$, we get:

$$\Pr(b \mid \mathcal{R}_2) \cdot \Pr(t_k, t_d \mid \mathcal{R}_2) \propto e^{-(1+2\lambda)b} \cdot \Pr(t_k, t_d \mid \mathcal{R}_2)$$

$$\Pr(b \mid \mathcal{R}_2) \propto e^{-(1+2\lambda)b}$$

Therefore,

$$\boxed{\mathbb{E}[b \mid \mathcal{R}_2] = \frac{\displaystyle\int_0^\infty be^{-(1+2\lambda)b}\,db}{\displaystyle\int_0^\infty e^{-(1+2\lambda)b}\,db} = \frac{1}{1+2\lambda}} \tag{99}$$

Since $\mathbb{E}[b|\mathcal{R}_1] = \mathbb{E}[b|\mathcal{R}_2]$, the solution is the same in both regimes, and no weights are needed. Therefore:

$$\boxed{\mathbb{E}[b|\mathcal{E}_8] = \frac{1}{1+2\lambda}} \tag{100}$$
